# FCD-11: A First-in-Class COMPASS Inhibitor in Cancer Therapeutics

**DOI:** 10.64898/2026.07.30.741768

**Authors:** Zibo Zhao, Meenal Karunanidhi, Sarah Gold, Luke St John, Rukkia Liaqat, Marta Iwanaszko, Claudia Bigas, Eita Uchida, Kentaro Mineji, Jun Watanabe, Salvador Aznar Benitah, Rintaro Hashizume, Rama K. Mishra, Ali Shilatifard

## Abstract

COMPASS (Complex of proteins associated with Set1) are highly conserved chromatin regulatory complexes responsible for all methylation marks on histone H3 lysine 4 (H3K4). The COMPASS protein SET1A is upregulated in metastatic breast cancer, and its H3K4 methyltransferase activity promotes metastasis in a palmitic acid diet setting. We identify and characterize FCD-11, a first-in-class small molecule inhibitor of COMPASS activity designed to disrupt the termolecular interface between COMPASS SET domains, ASH2L, and RBBP5. FCD-11 significantly inhibited the H3K4me3 methyltransferase activity of SET1A/COMPASS and MLL1/COMPASS in vitro. ChIP-seq and CETSA indicated that FCD-11 selectively inhibits SET1A/COMPASS activity in mouse embryonic stem cells and breast cancer cell lines. FCD-11 significantly reduced tumor size and extended survival in mouse models of breast cancer. Our findings establish FCD-11 as a potent, COMPASS-specific inhibitor lead compound with preclinical efficacy in breast cancer models and therapeutic potential for cancers with abnormal dependence on SET1A/COMPASS activity.

## Introduction

Histone methyltransferases (HMTases) are critical chromatin regulators and transcriptional co-activators that have been extensively investigated in the context of cancer and other diseases. So far, therapeutic development of HMTase inhibitors has been primarily limited to the HMTase targets that are most frequently mutated in cancer, such as MLL1 (*KMT2A*), which is affected by 11q:23 chromosomal translocation in the majority of infant and secondary acute myeloid leukemia (AML) (*1*). However, MLL1 is only one of six HMTases in the human COMPASS (Complex of proteins associated with Set1) family (*2*). COMPASS HMTases serve as the catalytic subunits of conserved chromatin regulatory complexes that deposit methylation marks on histone 3 lysine 4 (H3K4), in close association with transcriptional activation (*3*). The human COMPASS family has three functional branches: SET1A and SET1B are the primary depositors of H3K4 di- and tri-methylation (H3K4me2/me3) genome-wide (*4, 5*), establishing broad H3K4me3 peaks (*6*), MLL1 and MLL2 methylate H3K4 at developmental genes and bivalent promoters (*2*), and MLL3 and MLL4 monomethylate H3K4 at enhancers to regulate gene expression (*7, 8*). Each of these methyltransferases requires the presence of four core subunits (WDR5, RBBP5, ASH2L, and DPY30, collectively referred to as WRAD) within the COMPASS complex for their enzymatic activity (*9*). Other branch-specific COMPASS subunits are thought to contribute to complex stabilization and chromatin recruitment.

SET1A and SET1B are not often found to be mutated in cancer (*10*). However, several lines of evidence point to SET1A as a potentially impactful therapeutic target. SET1A is upregulated in metastatic breast cancer cell lines and may promote metastasis by increasing matrix metalloproteinase activity, as the loss of SET1A reduced metastasis in breast cancer mouse models (*11*). SET1A also regulates genomic stability by methylating H3K4 at stalled replication forks, which facilitates the mobilization of histones by FANCD2 (*12*). Additionally, SET1A activity is associated with metastasis in the setting of a palmitic acid diet (*13*), and SET1A depletion reduced lymph node metastasis in this setting (*13*). Together, these findings suggest that an inhibitor targeting SET1A/COMPASS activity could be translated as a therapeutic approach to limit tumor progression and metastasis in cancer patients.

We reasoned that COMPASS activity could be potentially targeted via a small molecule inhibitor with a complex-disrupting mechanisms of action. For example, the MLL1/COMPASS inhibitor revumenib disrupts the interaction between N-terminal MLL1 and the MLL1/2 branch-specific subunit Menin, evicting MLL1 oncofusion proteins from chromatin. This kind of N-terminal interaction disruptor is required for MLL1-rearranged cancers, in which the MLL1 C terminal (and MLL1 catalytic activity) is lost. However, for cancers with high expression of catalytically active COMPASS (such as SET1A/COMPASS in metastatic breast cancer), inhibiting catalytic activity would be the therapeutic goal. We therefore sought to develop small molecule inhibitors that could disrupt the catalytic activity-enabling interactions between the C-terminal domains of COMPASS HMTases and the subunits of their core WRAD module. Recent advances in our understanding of COMPASS complex structure and nucleosome binding have made the development of such an inhibitor possible (*14–16*).

Here, we report the development of a first-in-class small molecule COMPASS HMTase inhibitor. We describe optimization of in vitro HMTase assays that enabled us to identify more specific COMPASS inhibitors following an in silico screen using the available cryo-EM structure of nucleosome-engaged MLL1/COMPASS. We present data demonstrating that our lead compound FCD-11 specifically inhibits the H3K4me3 activity of SET1A/COMPASS and MLL1/COMPASS in vitro. We then demonstrate FCD-11 specificity and selectivity for SET1A/COMPASS in cells. We show that the effects of FCD-11 treatment resemble those of SET1A loss-of-function in murine embryonic stem cells (mESCs), and that FCD-11 treatment displays potent cytotoxic anti-tumor efficacy in SET1A-dependent breast cancer cell lines, where it also diminishes the broad H3K4me3 peaks deposited by SET1A/COMPASS. Finally, we demonstrate that administration of FCD-11 significantly reduces tumor burden and increases survival in mice with tumors established from orthotopic xenografts of different breast cancer cell lines. Our study provides molecular insight into the impact of SET1A/COMPASS inhibition in multiple cell lines and animal models of breast cancer, indicating a potential therapeutic avenue for the treatment of human cancers with abnormal SET1A/COMPASS activity.

## Results

### COMPASS HMTases prefer distinct substrates *in vitro*

Our approach to identification of COMPASS inhibitors began with in silico virtual screening. In order to evaluate candidates and develop a structure-activity relationship, we would require robust H3K4 methylation assays for biochemical screens. To establish optimal conditions for these assays, we systematically compared the substrate preferences of the six COMPASS HMTases using multiple recombinant substrates of increasing chromatin complexity. First, we evaluated the total HMTase activity of recombinant SET1A, SET1B, and MLL1-4 SET domain complexes obtained from Active Motif (fig. S1A) on histone H3 peptide (amino acids 1-25), free histone H3.1, and H3.1-containing mononucleosomes via MTase-GLO^TM^ assays (Fig. 1, A to C). These experiments revealed marked differences in substrate preference among COMPASS family members *in vitro*. SET1A only displayed strong activity toward free histone H3.1, with measurable activity towards nucleosomes that was measurable but significantly lower than that of all other COMPASS HMTases. In contrast, the closely related SET1B preferentially methylated histone H3.1 and histone peptides. MLL1-4 exhibited activities towards all three substrates, with the most significant MLL1 activity towards mononucleosomes (Fig. 1, A to C). Substrate preference was also observed for other HMTases that we would use for counterscreens: SETDB1, PRDM9, and G9a exhibited higher activity toward free histones and histone peptides, whereas DOT1L retained activity only on mononucleosome substrates (fig. S1B).

**Fig. 1.**
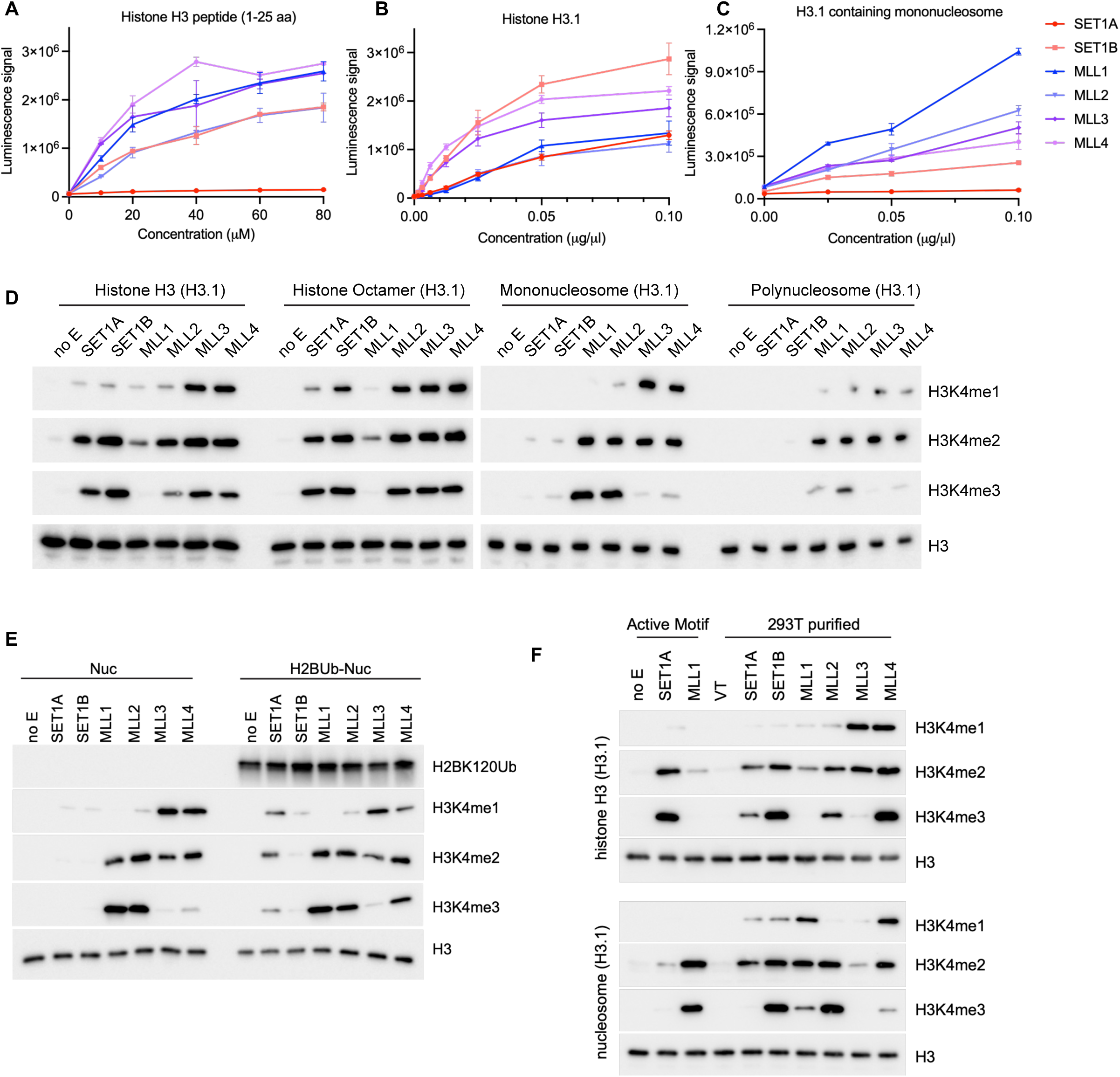
COMPASS methyltransferases prefer distinct substrates in vitro. (**A-C**) Analysis of in vitro methyltransferase activities for COMPASS complexes purchased from Active Motif. SET1A, SET1B and MLL1-4 SET complexes were evaluated via the MTase-GLO^TM^ assay using three different substrates: histone H3 peptide (1-25 a.a.) (**A**), histone H3.1 (**B**), and H3.1-containing mononucleosomes (**C**). Enzyme concentrations were optimized individually for each assay. n=3 for histone H3.1, n=2 for histone peptides and mononucleosomes. (**D**) Western blot comparison of in vitro H3K4 methyltransferase activity in assays performed with H3.1-containing histone H3, histone octamer, mononucleosome, or polynucleosome substrates. (**E**) Comparison of in vitro H3K4 methyltransferase activity in assays using unmodified mononucleosome or H2BK120Ub1 modified mononucleosome substrates. For all reactions, samples were incubated at room temperature for 2.5 hours. (**F**) Western blot comparison of in vitro H3K4 methyltransferase assays using COMPASS complexes purchased from Active Motif, versus those that were purified in-house. For in-house purification, 293T cells infected with lentivirus expressing 3xFlag tagged SET domain of SET1A, SET1B, or MLL1-4 were harvested, lysed, and immunoprecipitated against M2 slurry beads.

We then probed the effect of different substrates on differentiated methylation states of HMTase activity (H3K4me1/2/3), expanding our assays to include free histone H3.1, histone H3.1 octamers, histone H3.1-containing mononucleosomes, and histone H3.1-containing polynucleosomes (Fig. 1D). HMTase activity in these assays was evaluated via western blot and was consistent with the initial substrate comparison for total HMTase activity. SET1A and SET1B complexes exhibited maximal activity on free histone H3.1 and histone octamers, whereas MLL1 demonstrated maximal activity on mononucleosomes. MLL2 methylated free histone H3.1, histone octamer, and mononucleosomal substrates with similar efficiency. Finally, MLL3 and MLL4 displayed a slight preference for free histone H3.1 over histone octamers, with markedly reduced H3K4me3 activity on nucleosomal substrates (Fig. 1D). Similar substrate preferences were also observed using H3.3-containing substrates, indicating that histone variant composition does not substantially alter COMPASS enzymatic activity under these assay conditions (fig. S1C). Previous studies have reported that COMPASS activity can be directly stimulated by H2B ubiquitination, supported by structural evidence showing that the ubiquitin moiety primarily contacts the Set1 catalytic domain itself (*17–20*). Specifically, the Set1 arginine-rich motif (ARM) helix autoinhibits COMPASS upon nucleosome binding, whereas H2Bub allosterically activates COMPASS by anchoring the Set1 ARM helix. However, these studies were largely performed in yeast, where Set1 is most closely related to human SET1A/SET1B. Importantly, the unique ARM element is present only in Set1/SET1A/SET1B complexes and is absent from other COMPASS family HMTases. In human MLL1 complexes, H2B ubiquitination functions as a structural anchor that stabilizes the “active” conformation of MLL1 on the nucleosome through direct interaction between RbBP5 and ubiquitinated H2B (*14, 21*). In the absence of H2B ubiquitination, MLL1 docking on the nucleosome represents a lower-activity state rather than a completely inactive state (*21*). H3K4me3 has also been shown to be retained in the absence of H2B ubiquitination during myogenic differentiation (*22*). However, we also noticed conflicting observations in vitro. One study reported only minimal H3K4me3 activity for the MLL1 activity on in vitro reconstituted ubiquitinated nucleosome core particles (ubNCPs) (*14*). In contrast, another study demonstrated that MLL1 could efficiently catalyze H3K4me3 on unmodified nucleosome core particles, with substantially more prominent H3K4me3 signals (*23*). Based on these somewhat conflicting observations, we further examined the influence of H2B ubiquitination on MLL1 activity in our own experimental systems. Our results showed that H2BK120Ub1 modestly enhanced SET1A activity and shifted MLL4’s product distribution from mono-toward tri-methylation, with little impact on other COMPASS complexes under our assay conditions (Fig. 1E).

We were also concerned that the recombinant SET1A and SET1B complexes available from Active Motif lacked the DPY30 subunit of the core WRAD module (the MLL1-4 complexes contained all four WRAD subunits: WDR5, RBBP5, ASH2L, and DPY30) (fig. S1A). To compare the presence and absence of DPY30 on SET1A/B COMPASS activity, we first evaluated recombinant SET1A and SET1B/COMPASS complexes available from Reaction Biology, which contained all four WRAD subunits. Unfortunately, these complexes showed no detectable H3K4 methyltransferase activity under our assay conditions (fig. S1D). Therefore, we performed inhouse purification in an attempt to obtain fully assembled COMPASS HMTase complexes. We expressed 3xFlag-tagged SET domains of SET1A, SET1B and MLL1-4 in 293T cells, purified each via M2 agarose immunoprecipitation, and confirmed subunit composition by silver staining (fig. S1E). Western blot analysis of co-immunoprecipitation demonstrated intact WRAD interactions in the 293T cell-derived complexes (fig. S1F). In vitro HMTase assays showed that their substrate preferences recapitulated those of the in vitro-assembled recombinant complexes from Active Motif: preferential methylation of histone H3 or comparable activity toward histone H3 and mononucleosomes, with the exception of MLL1, which showed a clear preference for mononucleosome substrates (Fig. 1F).

Together, these assay optimization data defined the optimal substrates and conditions for assaying COMPASS-mediated H3K4 methylation in vitro. Based on the results, we decided to use unmodified H3.1-containing mononucleosomes as substrates for MLL1 and free histone H3.1 for all other COMPASS HMTases. Because DPY30 is not part of the ternary interface observed in the cryo-EM structure of the MLL1–RBBP5–ASH2L complex, we chose to use commercially available SET domain complexes from Active Motif over our in-house purifications to maintain consistency across assays.

### FCD-11 is a first-in-class COMPASS inhibitor

A structure of human MLL1 recognizing the nucleosome was recently resolved by cryo-electron microscopy (cryo-EM) to near-atomic resolution (3.2 Å) (*14*). This structure, which captured a COMPASS catalytic module binding the nucleosome for the first time, otherwise resembles cryo-EM and crystal structures of yeast COMPASS complexes as reported by our group and others (*15, 16, 24*). We decided to use the structure as a resource to identify potential inhibitors of COMPASS HMTase activity (COMPASS inhibitors) based on their potential to disrupt critical interactions between COMPASS subunits. We began by identifying the termolecular interface between the MLL1 SET domain, RBBP5, and ASH2L from the MLL1-ubNCP complex (PDB:6KIU) cryo-EM structure (*14*). We initially applied a site-finding algorithm to this structure and identified a potential small molecule binding pocket at the interface between the MLL1 SET domain and ASH2L. However, this binding pocket was blocked by an extended labile loop in RBBP5 (Fig. 2A). Therefore, we decided to carry out 100 ns molecular dynamics simulations (MDS) to generate a stable structural conformation of the termolecular complex. When we applied the site-finding algorithm to this MDS-generated structure, we observed an unobstructed small molecule binding pocket at the termolecular interface. MDS parameters and the residues in the putative binding pocket are detailed in the Methods section. Considering the potential binding pocket, we then performed *in silico* virtual high-throughput screening (vHTS) to search for potential COMPASS inhibitors that could disrupt the termolecular interface between the MLL1 SET domain, ASH2L, and RBBP5. The vHTS screening strategy is detailed in the Methods section. Our *in silico* screen generated 134 hits with a Glide extra precision score < - 6.0. Out of these hits, 99 compounds were selected for subsequent biochemical screening based on availability, pricing, and synthetic feasibility factors.

**Fig. 2.**
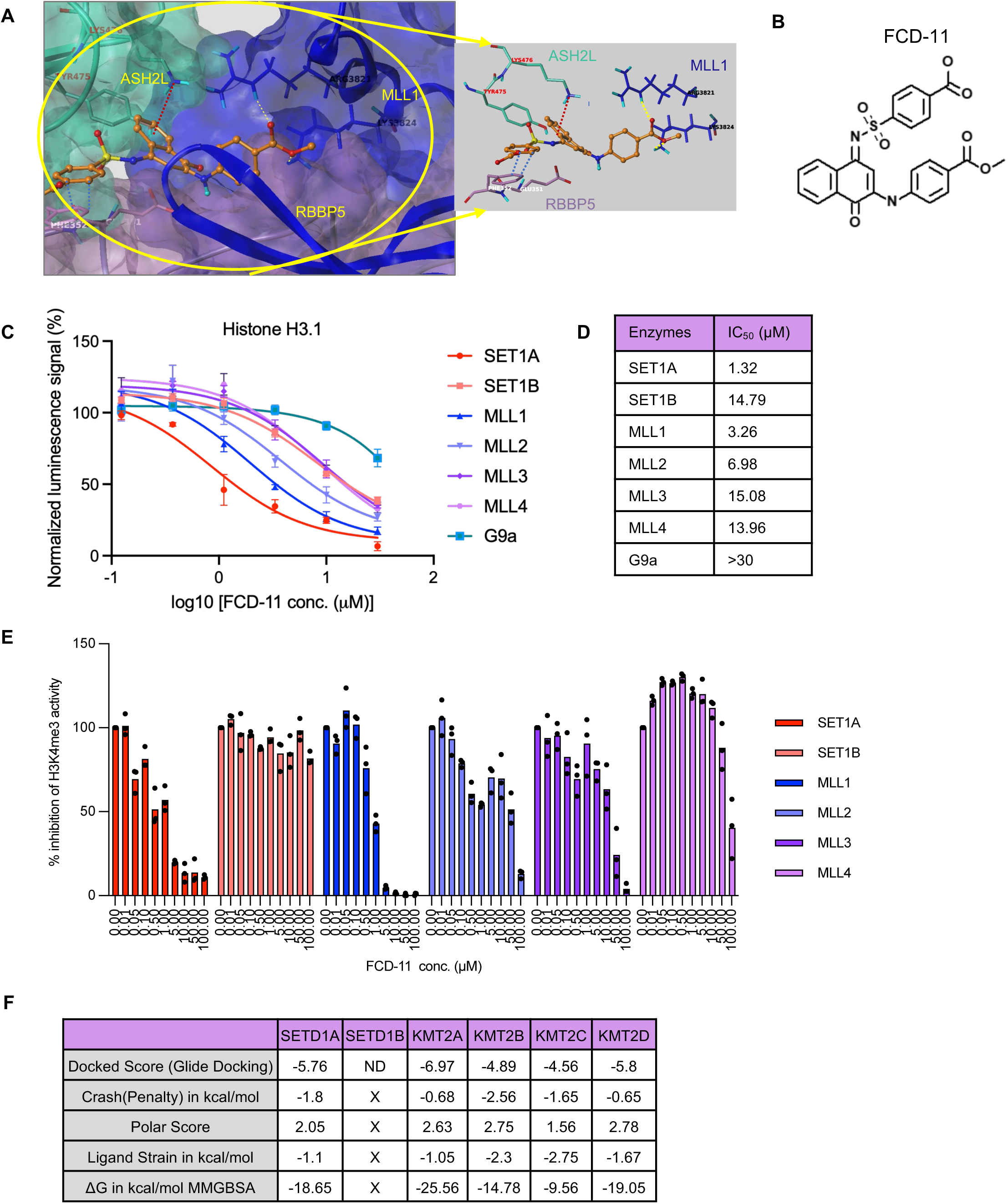
FCD-11 is a first-in-class COMPASS inhibitor. (**A**) 3D docked pose of FCD-11 (ball-and-stick model, orange) was built upon the termolecular MLL1 (blue), RBBP5 (light pink) and ASH2L (light green) complex interface (PDB ID: 6KIU). Yellow dotted lines indicate potential hydrogen bonds, light blue dotted lines indicate potential π-π interaction, and brown dotted line indicates potential π-cationic interaction. (**B**) Chemical structure of FCD-11. (**C-D**) FCD-11 inhibition of methyltransferases activity in the MTase-Glo^TM^ assays using histone H3.1 as substrates (n=4). FCD-11 inhibition curve (**C**) and absolute IC_50_ values (**D**) are shown (n=4). G9a is included as a negative control. (**E**) Bar plots showing the % inhibition of H3K4me3 activity in the presence of indicated concentrations of FCD-11 for COMPASS HMTase complexes and the non-COMPASS HMTase G9a. Histone H3 was used as the substrate for all reactions except for those with MLL1, where mononucleosomes were used. Inhibition was analyzed via western blot for H3K4me3 (see Fig. S3E), performed in triplicates, with band intensity quantified by ImageJ. (**F**) Docked Score of FCD-11 on COMPASS methyltransferases. AlphaFold Thread of Phyre2 Homology Model building tools were used to generate structures of all the COMPASS family members and then replaced MLL1 from cryo-EM structure by these newly built SETs structures.

Through two rounds of SET1A/COMPASS H3K4me3 methyltransferase inhibition assays (fig. S2, A to C) and a counter screen with G9a H3K9me2 methyltransferase inhibition assays (fig. S2D), we identified the compounds M2, M58, and M62 (fig. S2E) as inhibitors of SET1A/COMPASS activity. Next, we used the structures of these three compounds to generate a 5-point common pharmacophore query, which we then used to screen a curated library for potential SET1A/COMPASS inhibitors. The pharmacophore query generated 10 analogues that were very similar to the initial hits. Out of these 10 analogues, FCD-11 exhibited the most promising activity in our SET1A/COMPASS H3K4me3 methyltransferase inhibition assay (fig. S2F) (data for other analogues not shown), resulting in the identification of FCD-11 as a potential first-in-class SET1A/COMPASS inhibitor (Fig. 2B). The docked pose of FCD-11 at the termolecular interface of MLL1-ASH2L-RBBP5 revealed potential interactions with all three subunits, including classical H-bonds formed with the R3821 and K3824 residues of MLL1, a π-π interaction with F352 and a long-range electrostatic interaction with E351 of RBBP5, as well as a π-cationic interaction with K476 and a hydrophobic interaction with Y475 of ASH2L (Fig. 2A).

To assess the inhibitory effects of FCD-11 against COMPASS family HMTases and other HMTases, we titrated FCD-11 against SET1A, SET1B and MLL1-MLL4/COMPASS using our optimized HMTase inhibition assays (Fig. 1 and fig. S1), including both the MTase-Glo^TM^ high-throughput assay and in vitro HMTase assay followed by the western blot analysis of H3K4me1/2/3 with total H3 as loading control. Using free histone H3 as the substrate, FCD-11 showed the strongest inhibition toward SET1A, with an IC_50_ of 1.32 µM, followed by MLL1 with an IC_50_ of 3.26 µM (Fig. 2, C and D). Using histone H3 peptides as substrates, we observed similar selectivity toward SET1A and MLL1, while G9a, PRDM9, and SETDB1 served as negative controls and showed minimal sensitivity to FCD-11 (fig. S3, A and B). In assays using mononucleosome substrates, FCD-11 exhibited the strongest inhibition toward MLL1, whereas DOT1L was largely unaffected and served as a negative control (fig. S3, C and D). SET1A and SET1B could not be reliably evaluated in the MTase-Glo^TM^ assay under these conditions because they exhibited minimal enzymatic activity on unmodified mononucleosome substrates (Fig. 1B). However, using H2BK120 monoubiquitinated mononucleosomes, we observed a similar IC_50_ for FCD-11 against SET1A (fig. S3E), further supporting the robustness and substrate-independent inhibitory activity of FCD-11 toward SET1A/COMPASS. FCD-11 also exhibited comparable inhibition of MLL1 under both conditions (fig. S3F).

As an orthogonal validation approach, in vitro methylation assays followed by western blot analysis similarly demonstrated that SET1A was the COMPASS complex most sensitive to FCD-11 inhibition, consistent with the MTase-Glo^TM^ assay quantification (fig. S3G). Quantification of the in vitro methylation assays further demonstrated that FCD-11 most potently inhibited SET1A/COMPASS-mediated H3K4me3 activity, with an IC_50_ of approximately 0.46 µM (Fig. 2E). In contrast, SET1B and MLL3 activities were only weakly affected by FCD-11. FCD-11 also inhibited MLL4-mediated H3K4me1 activity and MLL1-mediated H3K4me3 activity, with an IC_50_ of approximately 0.85 µM for MLL1. Together, these results demonstrate the selective inhibitory activity of FCD-11 toward specific COMPASS HMTases.

We could not directly identify candidates with likely specificity for SET1A/COMPASS via our *in silico* approach because the cryo-EM structures of MLL1 or MLL3 recognizing the nucleosome were the only relevant published structures currently available (*14*). In the in vitro HMTase assays, we observed that FCD-11 inhibited not only SET1A activity, but also the activities of MLL1 and MLL4. Therefore, to structurally characterize FCD-11 engagement with different COMPASS family members, we used AlphaFold Thread of Phyre2 Homology Model building tools to generate structures of SET domains for all the COMPASS family HMTases and then used these to replace MLL1 in the existing cryo-EM structure. After building and validating the complex of these SET domains with ASH2L and RBBP5, we then docked FCD-11 and finally computed binding energy scores (ΔG) for FCD-11 to all COMPASS family members (Fig. 2F). Consistent with the pattern of inhibition in the in vitro HMTase activity assay, the lowest ΔG for FCD-11 was seen with MLL1, SET1A, and MLL4 (Fig. 2F).

To further explore the structure-activity relationship of FCD-11, we first identified 18 analogues through matched molecular pair analysis using the CheEMBL database (*25*) with FCD-11 as the query. Next, we evaluated the FCD-11 analogues in H3K4me3 methyltransferase inhibition assays (fig. S4A). After validation at lower concentrations, three compounds with IC_50_ values similar to FCD-11 were identified: Q15, Q16, and Q17 (fig. S4, B to D). Together, these results indicate that FCD-11 disrupts the ASH2L-RBBP5 interaction, and that this disruptive activity is correlated with SET1A/COMPASS HMTase inhibition.

### FCD-11 treatment resembles genetic loss of catalytic SET1A/COMPASS activity in mESCs

FCD-11 specifically inhibited the HMTase activity of SET1A and MLL1 in our in vitro assays (Fig. 2C and fig. S3, A to D), suggesting selectivity among COMPASS family HMTases. We therefore hypothesized that FCD-11 could selectively inhibit the H3K4me3 methyltransferase activity of these COMPASS HMTases in living cells. To further investigate the specificity and selectivity of FCD-11, we established a series of isogenic mESC (murine embryonic stem cell) lines harboring combinations of mutation or knockout (KO) of COMPASS HMTase including SET1AΔSET, SET1BKO, MLL1KO, MLL2KO, SET1AΔSET / SET1BKO (ABKO), SET1AΔSET / SET1BKO / MLL1KO (AB1KO), SET1AΔSET / SET1BKO / MLL2ΔSET (AB2KO), MLL3KO, MLL4KO, and MLL3KO / MLL4KO (MLL34KO) lines (fig. S5A) (*6, 26–28*). Complete SET1AKO mESCs are not viable (*27*), so we used SET1AΔSET lines where the catalytic SET domain is deleted. First, we confirmed the overall H3K4 methylation phenotype in these lines. Bulk H3K4me3 levels were most significantly downregulated in the AB2KO cells, and to a lesser extent in MLL2KO cells, but were only minimally affected in SET1AΔSET cells, consistent with our previous study showing MLL2’s role in regulating bulk H3K4me3 levels in the absence of SET1A (fig. S5A) (*6*). Next, we performed RNA-seq to confirm the overall gene expression phenotype in our isogenic cell lines, finding that loss of either MLL2 or combined loss of two or three COMPASS HMTases (ABKO, AB1KO or AB2KO) had the most profound effect on gene expression change, whereas loss of SET1A catalytic function (SET1AΔSET) or MLL1KO alone had minimal effect (fig. S5, B and C). These results align with the bulk methylation results and those of our previous study showing that the catalytic activity of SET1A is dispensable for mESC self-renewal and proliferation (*6*). Finally, we evaluated the effect of FCD-11 treatment on the overall H3K4 methylation phenotype in WT mESCs, finding that FCD-11 treatment did not affect bulk levels of H3K4me1/2/3 or other histone H3 methylation marks (fig. S5D).

After these validation steps, we sought to determine the effect of FCD-11 treatment on COMPASS-specific patterns of H3K4me3 via ChIP-seq (fig. S5E). Our group previously showed that while SET1A and SET1B implement broad H3K4me3 peaks at highly expressed genes, MLL2 is responsible for narrow H3K4me3 peaks at less transcriptionally active promoters (*6*). This distinction makes it possible to investigate selective effects of FCD-11 treatment on COMPASS HMTase activity based on genome-wide H3K4me3 patterns and patterns at specific loci. Since the chromatin binding patterns of SET1A, MLL1, and MLL2 are all highly promoter specific (fig. S5F), we performed K-means clustering of H3K4me3 peaks at the promoters of protein-coding genes in WT cells (Fig. 3A). Our H3K4me3 ChIP-seq data revealed that 2 µM FCD-11 treatment in WT mESCs resembled the H3K4me3 phenotype in SET1AΔSET mESCs, with genome-wide diminishment of broad H3K4me3 peaks in either condition relative to untreated WT mESCs (Fig. 3A). Zooming in on specific cluster1 loci gave us further insight into the locus-specific efficacy of FCD-11 treatment. FCD-11 treatment decreased broad H3K4me3 at the promoter regions of *Xpo7*, *Zfp219* and *Slc39a7/Rxrb*, which are also occupied by higher levels of SET1A compared to MLL1 and MLL2 (Fig. 3, B to D). The decrease observed with FCD-11 is more modest than that seen with SET1A SET domain deletion. This difference likely reflects the fact that the two conditions, SET1AΔSET cells and short-term (48-hour) FCD-11 treatment, are inherently distinct in both timescale and mode of perturbation (genetic versus pharmacological). Compared to its analogues Q16 and Q17, FCD-11 exhibited the strongest effect in diminishing H3K4me3 signal at promoter regions in cluster1 (fig. S5G). Consistent with the bulk level changes of histone H3 methylation status with FCD-11 treatment, other major methylation marks including H3K27me3, H3K36me2/3, H3K9me2/3 and H3K4me1/2 were not significantly influenced by FCD-11 treatment (fig. S5H).

**Fig. 3.**
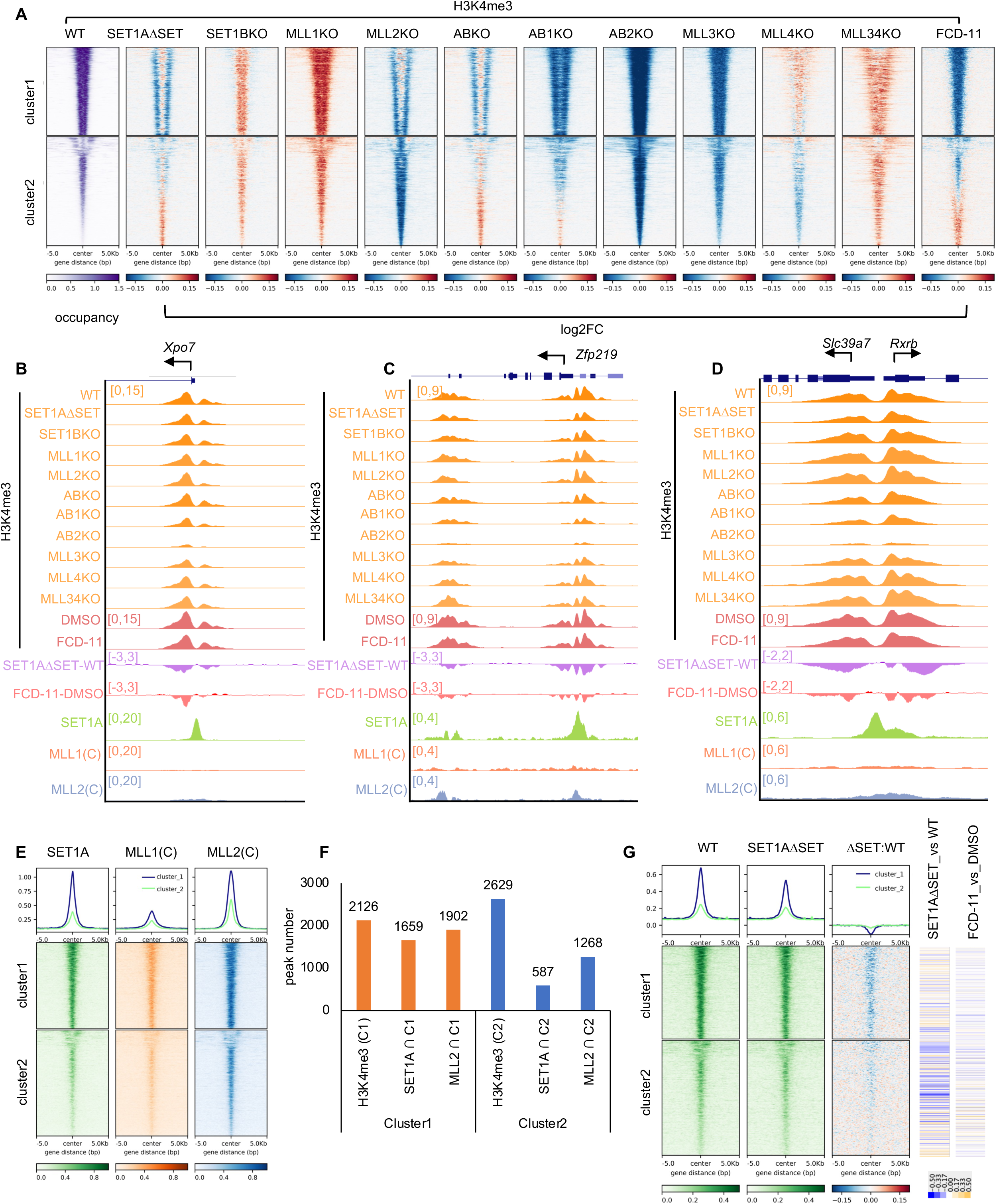
FCD-11 treatment resembles functional SET1A/COMPASS loss in mESCs. (**A**) H3K4me3 ChIP-seq was performed in the indicated mESC cell lines with spike-in control. Heatmaps of log2 fold change in the signal intensity of TSS-proximal H3K4me3 peaks (n=4756) in COMPASS KO and/or ΔSET relative to WT mESCs. Peak centers are sorted in descending order of H3K4me3 peak width and two peak clusters are shown: broad (cluster1) and narrow (cluster2) peaks. The catalytic SET domain of SET1A has been deleted in the SET1AΔSET mESCs. ABKO = SET1A/SET1B KO; AB1KO = SET1A/SET1B/MLL1 KO; AB2KO = SET1A/SET1B/MLL2 KO; MLL34KO = MLL3/MLL4 KO. For compound treatment, cells were treated with 2 µM FCD-11 for 48 hours before ChIP-seq experiments. (**B-D**) Track examples showing the H3K4me3 signals at the *Xpo7* (**B**), *Zfp219* (**C**) and *Slc39a7*/*Rxrb* (**D**) gene loci. Subtracted signals are shown SET1AΔSET versus WT and for FCD-11 treatment (2 µM for 48 hours) versus DMSO. (**E**) Metaplots and heatmaps of SET1A, MLL1 and MLL2 ChIP-seq occupancy for the two peak center clusters shown in (**A**). SET1A, MLL1 and MLL2 ChIP-seq were obtained from GSE200120 (*28*). (**F**) Quantification of H3K4me3, SET1A and MLL2 peaks in the two clusters. (**G**) Metaplots and heatmaps of SET1A ChIP-seq occupancy in WT and in SET1AΔSET cells, as well as log2FC of occupancy in SET1AΔSET versus WT, for the two peak center clusters shown in (**A**). At right, log2FC heatmaps display differential expression of genes near H3K4me3 peaks (in both cluster1 and cluster2) for SET1AΔSET versus WT, and for FCD-11 versus DMSO.

When we centered SET1A, MLL1, and MLL2 ChIP-seq signals onto H3K4me3 peaks in WT mESCs, we observed that SET1A appears to preferentially occupy regions with broad H3K4me3 peaks (Fig. 3, E and F), consistent with our H3K4me3 ChIP-seq data in isogenic cell lines (Fig. 3A). Of note, we observed a different pattern of H3K4me3 decrease in SET1AΔSET cells compared to the FCD-11 treatment (Fig. 3A). Compared with the SET1AΔSET condition, the effect of FCD-11 treatment appears stronger in terms of the H3K4me3 reduction for cluster1. This may be explained by the fact that, despite their lack of SET domains, SET1A proteins are stably present on chromatin in SET1AΔSET cells (Fig. 3G). The shoulders of SET1A peaks also display a more profound H3K4me3 reduction than peak centers in SET1AΔSET relative to WT cells, suggesting that the SET domain-independent binding of SET1A might prevent demethylases from removing H3K4me3 at the same sites. We further examined gene expression changes associated with cluster 1 peaks, which are characterized by broad H3K4me3 peaks and high SET1A occupancy. Consistent with our earlier observations, transcriptional changes in these regions are relatively subtle between SET1AΔSET and WT, and FCD-11 treatment similarly induces minimal gene expression alterations (Fig. 3G and fig. S5, I and J). We note that the modest transcriptional effects observed in the SET1AΔSET condition are likely driven (at least in part) by indirect mechanisms, given that the gene expression changes clustered in regions with lower SET1A occupancy (Fig. 3G). Finally, given the essential role of SET1A’s catalytic activity during embryoid body (EB) differentiation (*27*), we tested whether FCD-11 mimics the effect of SET1AΔSET on EB formation from mESCs. FCD-11 treatment (2 µM or 5 µM) led to a reduction in EB size, partially resembling the phenotype observed with SET1AΔSET (fig. S5K). Taken together, these data clearly demonstrate that the COMPASS-specific inhibitory activity that we observed for FCD-11 in vitro is recapitulated in living cells treated with FCD-11, and suggest that FCD-11 is specific and selective for SET1A/COMPASS in the cellular context.

### FCD-11 treatment inhibits breast cancer cell viability, diminishes SET1A/COMPASS-deposited H3K4me3 peaks, and impairs in vivo tumor growth

After determining that FCD-11 acts as a selective SET1A/COMPASS inhibitor in mESCs, we sought to evaluate its anti-tumor potential. We began by submitting FCD-11 for NCI-60 cell line screening in a single-dose study at NCI DTP (Developmental Therapeutics Program) (fig. S6A). Overall, FCD-11 demonstrated cell growth inhibition activity in leukemia, melanoma, ovarian cancer, renal cancer, and breast cancer cell lines (fig. S6A). Based on these results and recent results in the literature indicating that SET1A is upregulated in metastatic breast cancer cell lines (*11*) and that the catalytic activity of SET1A is required for metastasis of palmitic acid-mediated squamous cell carcinoma (*13*), we also chose several squamous cell carcinoma, glioma, and breast cancer cell lines for cell-based viability assays to evaluate FCD-11 cytotoxicity. We found that FCD-11 was most effective against the MDA-MB-453 and MDA-MB-468 breast cancer lines (fig. S6B). Interestingly, SET1A/COMPASS is particularly essential for breast cancer cell viability as compared to other COMPASS (Fig. 4A), offering a unique opportunity to target SET1A’s activity using FCD-11. When we analyzed the relationship between SETD1A dependency and FCD-11 IC_50_ across all of the tested breast cancer cell lines, we observed a positive correlation between SETD1A dependency and FCD-11 IC_50_ (Fig. 4B), indicating that cells with greater dependence on SETD1A tend to exhibit increased sensitivity (lower IC_50_) to FCD-11. We further extended this analysis to other COMPASS HMTases, SET1A/B-COMPASS subunits, and additional non-COMPASS HMTases. Among these, the strongest positive correlations were observed for SET1A (Fig. 4B) and SET1B (fig. S6C), whereas other factors showed minimal association or even negative correlation. However, given our experimental evidence that SET1B is not functionally inhibited by FCD-11 (Fig. 2C), we conclude that FCD-11 retains its primary specificity toward SET1A in breast cancer cells. When we treated MDA-MB-453 and MDA-MB-468 breast cancer cells with increasing concentrations of FCD-11 and its analogues Q15, Q16, and Q17, we observed that FCD-11 inhibited breast cancer cell viability most potently with IC_50_ ∼ 2.5 µM (Fig. 4C).

**Fig. 4.**
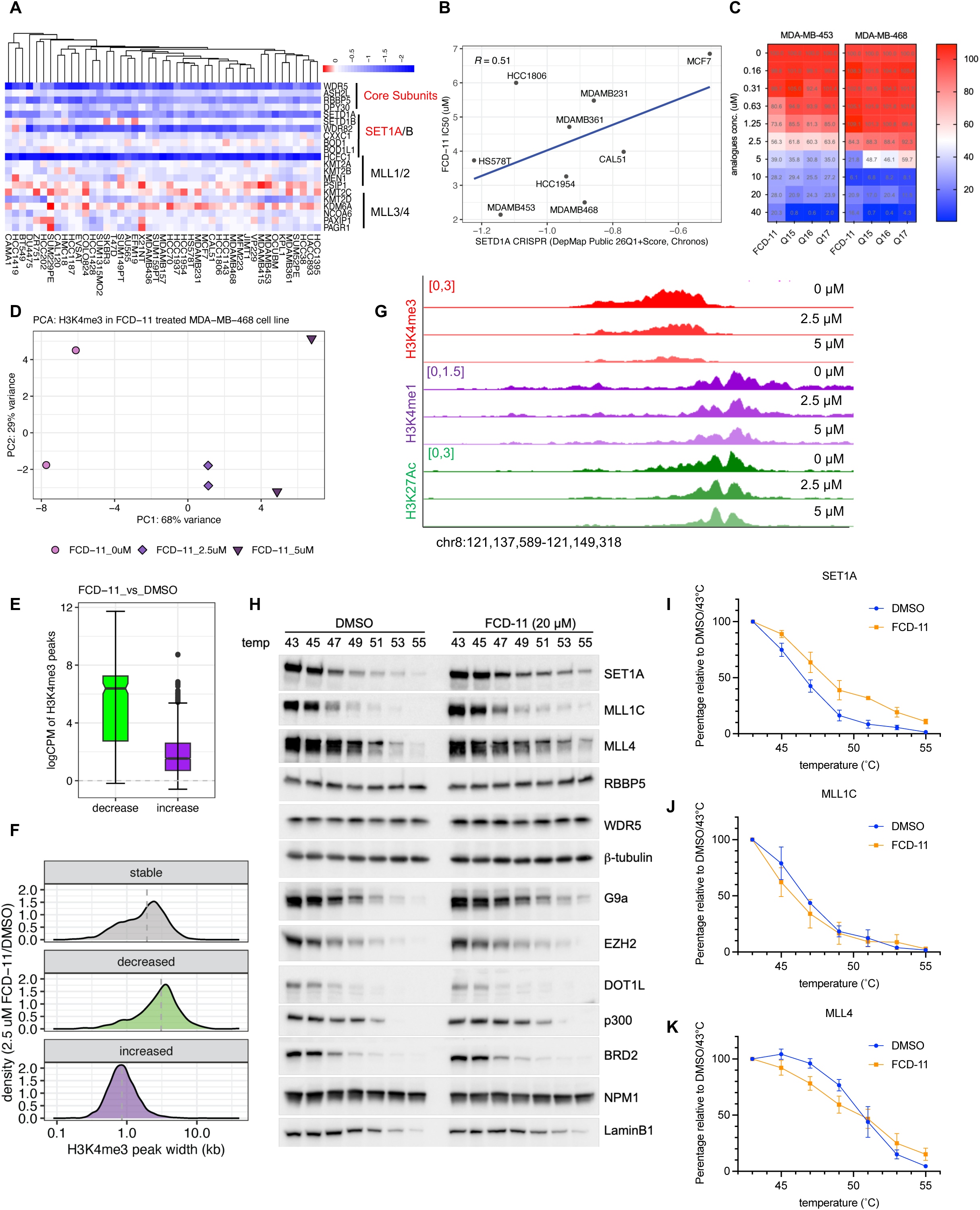
FCD-11 decreases cell viability, diminishes SET1A/COMPASS-deposited H3K4me3 peaks, and selectively engages SET1A/COMPASS in breast cancer cell lines. (**A**) Heatmap showing the CRISPR gene effect scores of COMPASS subunit knockouts in all CCLE breast cancer cell lines (DepMap Public 22Q2). (**B**) Correlation between SETD1A CRISPR score (SETD1A dependency score) and FCD-11 IC_50_ in the tested breast cancer cell lines. (**C**) CellTiter-Glo® luminescent cell viability assay of MDA-MB-453 and MDA-MB-468 cells treated with increasing concentrations of FCD-11 or its analogues Q15, Q16, or Q17 for 72 hours. Values shown as percentage of survival relative to DMSO. (**D**) Principal component analysis (PCA) of H3K4me3 ChIP-seq with DMSO (FCD-11 0 uM) or FCD-11 treatment at 2.5 µM or 5 µM for 48 hours. (**E**) Boxplot of logCPM (log counts per million) values for differential H3K4me3 peaks altered upon FCD-11 treatment at 2.5 µM relative to DMSO. Green, significantly downregulated peaks; purple, significantly upregulated peaks. adj.p < 0.01. (**F**) Density plot of peak width for the differential H3K4me3 peaks. (**G**) Track example showing representative alteration in H3K4me3 but not H3K4me1or H3K27Ac peaks upon FCD-11 treatment in MDA-MB-468 cells. All ChIP-seq samples are spike-in normalized. (**H**) Western blot analysis of the Cellular Thermal Shift Assay (CETSA) (*61*) investigating the stability of various COMPASS components (SET1A, MLL1C, MLL4, RBBP5, WDR5), other non-COMPASS methyltransferases (G9a, EZH2, and DOT1L), and non-relevant proteins (p300, BRD2, NPM1, and LaminB1) in MCF7 cells upon FCD-11 treatment. β-tubulin was used as a loading control. n=3. (**I-K**) Densitometry quantification of SET1A (**I**), MLL1C (**J**) and MLL4 (**K**) in the CETSA assay. n=3.

Next, we investigated the on-target efficacy of FCD-11 against the catalytic activity of SET1A in breast cancer cells. We treated cells with 0, 2.5, or 5 µM FCD-11 and performed differential ChIP-seq analysis of H3K4me3 peaks with spike-in normalization (Fig. 4D). Peaks that were significantly downregulated by FCD-11in breast cancer cells tended to have a greater peak width, consistent with SET1A’s role in depositing broad H3K4me3 marks genome-wide (Fig. 4, E to G and fig. S6D). We also observed activation of narrow H3K4me3 peaks in breast cancer cells (Fig. 4, E and F), consistent with the pattern in mESCs (Fig. 3A), though the impact of FCD-11 appeared to be more pronounced in breast cancer cells. This would be consistent with a dependence on SET1A that is not present in mESCs: for example, mESCs tolerate loss of SET1A catalytic activity via SET domain deletion. However, differences in culture conditions may also influence cellular responses to FCD-11.

To demonstrate on-target binding for FCD-11, we then applied a cellular thermal shift assay (CETSA) in MCF7 breast cancer cells to investigate the thermostability of various COMPASS subunits when treated with 20 µM FCD-11 for 3 hours. The thermostability of SET1A was significantly enhanced in the presence of FCD-11, indicating that FCD-11 binds to SET1A in these cells (Fig. 4, H and I), whereas the stabilization of MLL1 and MLL4 was not significant (Fig. 4, J and K). We also included other cellular proteins and found no significant alteration in their stability under the same conditions (Fig. 4H). Consistent with ChIP-seq data in mESCs and breast cancer cells, these CETSA results indicate that FCD-11 is specific and selective for SET1A/COMPASS in these cellular contexts.

Due to the essential role of SET1A/COMPASS (Fig. 4A) and the results of our cell viability assays (Fig. 4B) in breast cancer cell lines, we hypothesized that FCD-11 treatment would also suppress tumor growth in vivo. To determine the anti-tumor activity of FCD-11, we generated orthotopic xenograft tumors in nude mice via implantation of MDA-MB-468 (Fig. 5A), MDA-MB-453 (Fig. 5B), MDA-MB-231 (Fig. 5C) or HCC-1954 (Fig. 5D) human breast cancer cell lines into mammary fat pads and treated mice with FCD-11 (10 mg/kg) or vehicle control (DMSO) intraperitoneally when the tumor size reached 100 mm^3^. Mice were euthanized when the tumor size reached 1,000 mm^3^. FCD-11 treatment significantly inhibited mammary tumor growth and extended the survival of breast cancer xenograft recipient mice compared to the control treatment group in all four models tested (Fig. 5). Together, these data demonstrate anti-tumor efficacy and on-target activity for FCD-11 in breast cancer cells, indicating potential for future therapeutic development of FCD-11 or related compounds.

**Fig. 5.**
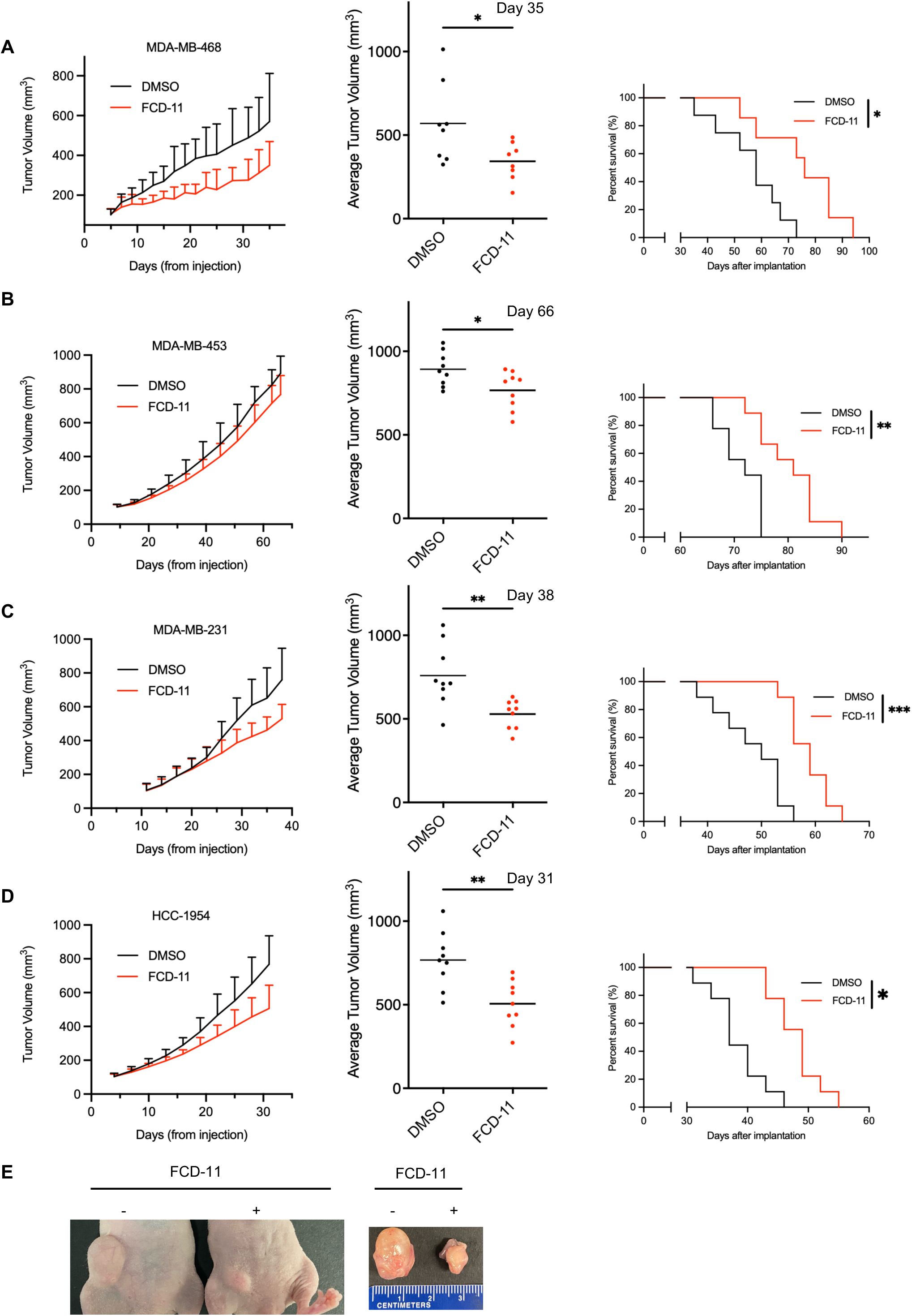
FCD-11 treatment impairs in vivo growth of breast cancer xenograft tumors. (**A-D**) Tumor development after injection of 4 × 10^6^ of MDA-MB-468 (**A**), MDA-MB-453 (**B**), MDA-MB-231 (**C**), or HCC-1954 (**D**) cells into the right mammary fat pads of nude mice. Following implantation, mice were treated with either vehicle (DMSO, n = 9) or FCD-11 (10 mg/kg, n = 9) daily for 7 days. Tumor growth was tracked up to the endpoint at which the tumor of a mouse in the control group reached maximum allowed size. Growth plots for mammary tumors in each treatment group are shown with mean tumor volume (mm^3^) and upper SD (left). Dot plots are shown for mammary tumor volume at growth tracking endpoint (center, days post-tumor cell injection indicated). Unpaired t-test was used for comparisons between DMSO and FCD-11 treatment. DMSO, n = 8, FCD-11 (10 mg/kg), n = 8. Animal survival was tracked for the remainder of the experiment, and survival curves are shown (right). Log-rank (Mantel-Cox) test was used for comparisons between DMSO and FCD-11 treatment. **e,** Photographs of nude mice inoculated with MDA-MB-468 cells and treated with vehicle or FCD-11 (left) and the mammary tumors taken from these mice (right).

## Discussion

In this study we report the compound FCD-11 (Fig. 2B), which we have validated as a first-in-class selective inhibitor of COMPASS activity *in vitro* (Fig. 2, C to E and fig. S3), with apparent specificity for SET1A COMPASS in isogenic mESC lines (Fig. 3A) and in human breast cancer cell lines (Fig. 4, B and C). Moreover, we have demonstrated anti-tumor efficacy for FCD-11 treatment in breast cancer cell lines (Fig. 4, A and B and fig. S6, A and B) and *in vivo* in orthotopic xenograft-derived mammary tumors (Fig. 5). FCD-11 is a drug-like analogue of three compounds that inhibited SET1A/COMPASS activity in our initial screen for potential COMPASS inhibitors (fig. S2-4). This screen (which is not the focus of the current report) also identified compounds that inhibit the HMTase activity of other COMPASS members via their corresponding HMTase assays; these will be discussed in future reports.

Our evaluation of substrates for COMPASS HMTase assays provides a reference point for future studies, including new information regarding substrate preference in vitro. For example, previous studies had demonstrated that MLL1 exhibited activities on both free histone H3 and nucleosome substrates in the absence of DPY30 (*29*), but that the presence of DPY30 completely abolished MLL1’s activity on free histone substrates and significantly boosted its activity on nucleosomes (*23*). Further studies had shown that the inclusion of DPY30 increased the activities of all COMPASS enzymes on nucleosome substrates by stabilizing the core WRAD module via its engagement with the intrinsically disordered regions of ASH2L (*30, 31*). However, to our knowledge no previous studies had directly compared the activities of SET1A and SET1B on histone versus nucleosome substrates in the absence of DPY30. We performed this comparison, and our results indicate that in the absence of DPY30, free histones or histone octamers serve as much better substrates for SET1A and SET1B than nucleosomes or polynucleosomes.

Besides the H3K4me3-depositing catalytic activity of SET1A, it is worth noting that catalytic-independent activities of SET1A and SET1B/COMPASS are also thought to contribute to tumorigenesis. We have shown that a cytoplasmic form of SET1B is essential for breast cancer cell survival independent of its SET domain (*32*). Others have shown that SET1A plays a critical role in DNA damage response through regulation of cyclin-K(*33*) and that it regulates mitochondrial respiration via transcriptional pause release of heme biosynthesis genes in acute myeloid leukemia(*34*). Additionally, it has been reported that SET1A can methylate non-histone substrates such as YAP in lung cancer (*35*).

Extensive preclinical investigation has previously supported clinical evaluation of selective inhibitors targeting catalytic activity of the HMTases DOT1L and EZH2 (which methylate H3K9 and H3K27, respectively). However, the potential for therapies targeting HMTases remains largely unrealized, and novel strategies (beyond substrate-competitive inhibitors) may be required to therapeutically target HMTases in cancer and other diseases (*1*). So far, the only HMTase inhibitor to have met with any clinical success is tazemetostat, which was FDA-approved for the treatment of certain patients with epithelial sarcoma or follicular lymphoma in 2020. Tazemetostat is a S-adenosyl-methionine (SAM)-competitive inhibitor of the PRC2 complex catalytic subunit EZH2 (all HMTases require SAM as a methyl donor cofactor). However, an emergent class of inhibitors have been developed to target EZH2 indirectly by disrupting its interaction with the PRC2 complex subunit EED (*36, 37*). In general, this kind of complex-disrupting inhibitor may represent a better way to target both the catalytic and catalytic-independent functions of HMTases (*1*). As discussed above, inhibitors have previously been developed to evict MLL1 oncofusion proteins from chromatin by disrupting interaction between the MLL1 portion of the oncofusion protein and the MLL1/2 branch-specific subunit Menin. More recently, inhibitors have also been developed to inhibit the catalytic activity of intact MLL1/COMPASS by disrupting the unique interaction between the N terminal of MLL1 and WDR5, a subunit of the core WRAD module (*38, 39*).

It should be emphasized that the initial screen leading to the identification of FCD-11 was performed to identify molecules that could disrupt the termolecular interactions between multiple subunits within the catalytic modules of COMPASS complexes, thereby enabling a complex-disrupting strategy rather than conventional substrate-competitive inhibition. This general COMPASS-disrupting approach allowed for identification of individual COMPASS HMTase-specific inhibitors via subsequent biochemical assays. A major limitation of the current study is that the basis of FCD-11 selectivity for SET1A (among other COMPASS) in cells is not yet clear, since our in vitro screen demonstrated that FCD-11 also inhibited MLL1’s H3K4me3 and MLL4’s H3K4me1 activities. Future studies will utilize isothermal calorimetry to quantitatively define the binding affinities of FCD-11 for ASH2L, RBBP5 and other WRAD subunits. By performing these studies with mutations at key ligand binding site residues, we can also assess the potential for resistance due to mutations at the target interface. Moreover, structural modeling predicts differences in binding pocket architecture among COMPASS HMTases (Fig. 2F) that could be leveraged to develop next-generation compounds with improved selectivity.

Our initial preclinical studies of FCD-11 have focused on its potential anti-tumor efficacy in breast cancer. However, SET1A/COMPASS has been reported to mediate cancer development and progression not only in breast cancer (*11, 40, 41*), but also in palmitic acid-induced metastatic oral carcinomas and melanomas (*13*), acute myeloid leukemia (AML) (*35*), colorectal cancer, and lung cancer (*35*). In line with these reports, our preliminary NCI-60 cell line screen demonstrated potential anti-tumor efficacy for FCD-11 in additional cancer types including leukemia, melanoma, and renal cancer (fig. S6A), and our cell viability assays further indicate cytotoxicity in squamous cell carcinoma and glioma cell lines (fig. S6B). Future studies may therefore explore broader therapeutic potential for FCD-11 across diverse cancer contexts. For breast cancer, future preclinical studies of FCD-11 would include more extensive animal models and/or patient-derived organoid models to determine subtype and genotype-specific anti-tumor efficacy. Additionally, while our current in vivo studies establish proof-of-concept antitumor efficacy, future studies to support further translational development will focus on defining PK/PD properties, measuring target engagement in tumor tissues, and integrating these parameters with efficacy outcomes, alongside safety and toxicity profiling. As a first-in-class COMPASS inhibitor, FCD-11 will serve as a lead compound for further optimization and development of COMPASS complex-disrupting cancer therapeutics.

## Materials and Methods

### Cell culture

MDA-MB-468, MDA-MB-453, and MCF7 cell lines were purchased from ATCC. CAL51 cells were purchased from DSMZ. SCC-25 cells were a kind gift from Dr. Salvador Benitah’s lab. SF8628 and KNS42 were kind gifts from Dr. Rintaro Hashizume’s lab. Hs578T, HCC1806, MDA-MB-231, HCC1954, MDA-MB-361, and BT474 cell lines were kind gifts from Dr. Maciej Lesniak’s lab. All above cells except SCC-25 were grown in Dulbecco’s modified Eagle’s medium (DMEM) with 10% FBS (Gibco). SCC-25 cells were grown in a 1:1 mixture of DMEM and Ham’s F12 medium containing 1.2 g/L sodium bicarbonate, 2.5 mM L-glutamine, 15 mM HEPES and 0.5 mM sodium pyruvate and supplemented with 400 ng/ml hydrocortisone and 10% FBS. V6.5 ESCs were grown in N2B27 medium supplemented with two inhibitors (2i) and LIF as described previously (*26*). For CRISPR knockouts and alterations of COMPASS HMTases, the sgRNA guide sequences have been reported in previous studies: SET1AΔSET (*27*); SET1BKO, MLL2ΔSET, MLL2KO (*6*); MLL1KO (*26*), MLL3/MLL4KO (*28*). The FCD-11 compound was custom synthesized by Princeton BioMolecular Research with > 90% purity.

### Antibodies and western blot

The following primary antibodies were used in this study: anti-H3K4me1 (Cell Signaling Technology (CST, #5326), anti-H3K4me2 (CST, #9725), anti-H3K4me3 (CST, #9751), anti-H3K27me3 (CST, #9733), anti-H3K9me2 (CST, #4658), anti-H3K9me3 (CST, #13969), anti-H3K27me2 (CST, #9728), anti-H3K27me3 (CST, #9733), anti-H3K27Ac (CST, *#*8173), anti-H3K36me2 (CST, #2901), anti-H3K36me3 (CST, #4909), anti-H3K79me1 (CST, #12522), anti-H3K79me2 (CST, #5427), anti-H3K79me3 (CST, #74073), anti-H3 (CST, #4499), anti-H2BK120Ub (CST, #5546), anti-MLL4 (NT, generated in-house), anti-MLL2 (CT, generated in-house), anti-SET1A (CT, generated in-house), anti-SET1B (NT, generated in-house), anti-MLL1C (CST, #14197), anti-RbBP5 (Fortis Life Sciences, A300-109A), ASH2L (CST, #5019), anti-WDR5 (CST, #13105), anti-DPY30 (Fortis Life Sciences, A304-296A), G9a (CST, #3306), EZH2 (CST, #5246), DOT1L (CST, #77087), p300 (CST, #86377), BRD2 (CST, #5848), NPM1 (CST, #3542), LaminB1 (Proteintech, 66095-1-Ig), Rpb1 NTD (CST, # 14958), Hsp90 (Santa Cruz Biotechnology, sc-13119), and anti-ß-tubulin (Proteintech, 66240-1-Ig). Secondary antibodies used in the study are mouse anti-rabbit IgG (Jackson immunoresearch, 211-032-171) and goat anti-mouse IgG (Jackson immunoresearch, 115-035-174). Western blot analysis was performed as previously described(*42*). For SET domain protein purification, the SET domains of SET1A (1450-1707aa), SET1B (1669-1966aa), MLL1 (3735-3972aa), MLL2 (2508-2715aa), MLL3 (4707-4911aa) and MLL4 (5337-5537aa) were custom synthesized by GenScript and cloned into the pLJM1 vector with 3xFlag tag. Lentiviruses expressing the SET domain proteins were infected into 293T cells for large scale purification using M2 agarose beads. Eluted samples were concentrated and validated with silver staining, western blot, and in vitro H3K4 HMTase assay.

### Molecular dynamics simulations (MDS) and binding pocket identification

The MLL1-ubNCP complex cryo-EM structure (PDB:6KIU) (*14*) was modified to correct missing atoms and side chains and undesired residue orientations (Asn, Gln, and His) using the Prime validation tool from the Schrodinger suite (*43*). The termolecular complex interface of the MLL1 SET domain, RBBP5, and ASH2L was considered for the MDS using the Desmond module implemented in Schrodinger platform. The termolecular complex was subjected to a protein preparation panel prior to preparation in a TIP3 water box for MD simulations. A 100 ns MDS was carried out with a time step of 100 ps in an isotropic pressure coupling condition. We also applied a smooth particle mesh Ewald method for accurate long range electrostatic evaluation in an NTP ensemble attaching a Nose-Hoover thermostat. The stability of the complex was determined by analyzing various output such as the plot of potential energy vs time, and RMSD deviations of the backbone and side chains. Considering this MDS-generated structure, a putative small molecule binding pocket was identified at the termolecular interface of MLL1, ASH2L and RBBP5 via implementation of the SITE-ID algorithm in the Schrodinger platform (*44*). The algorithm-identified binding pocket contained the residues R3821, K3824, E3857, I3920, D3921, Q3923, and H3925 from MLL1, E341, N342, V343, Y345, E351 and F352 from RBBP5, and E310, K311, G312, Y313, Y475, and K476 from ASH2L. Adjacent residues were also added to define the putative ligand binding pocket for vHTS.

### Virtual high-throughput screening (vHTS)

A curated library of ∼10 million drug-like compounds was prepared for vHTS by filtering the public ZINC database (*45*) (∼18 million compounds at time of study completion) to eliminate promiscuous, non-drug-like and toxic compounds (*43*). A 3-tier sequential screening algorithm was implemented as Glide in the Schrodinger suite (version: Schrodinger 2019-4). For this grid-based algorithm, a 12×12×12Å^3^ grid box was defined in the putative ligand binding pocket in consideration of all critical residues in the MLL1 SET domain, ASH2L, and RBBP5. The OPLS2005 force field was used, and the ligand van der Waals radii was set to 0.8Å with a partial atomic charge < 0.15 esu.

### PHYRE2.2 Alpla Thread Prediction

Alpha Thread is a recent implemented feature in PHYRE2.2 (*46*), a well-known homology modeling tool that relies on template structures for accurate prediction. After submitting the sequences of the other COMPASS family members, the Alpha Thread incorporated in PHYRE2.2 searched the AlphaFold Protein Structure Database (*47, 48*) using Blastp to find the initial structures for the other COMPASS family members. Then these template structures were used in PHYRE2.2 to optimize the side chains of the different residues. The final structures generated from Alpha Thread guided PHYRE2.2 were subjected to MolProbity validations (*49*) yielding scores above 90%. These validated structures were then used for molecular docking experiments with the FCD-11 compound across various COMPASS family members.

### H3K4 methyltransferase assay quantified by western blot

A 20 µL reaction containing recombinant methyltransferase or protein complex (0.1 µg for SET1A, Active Motif #81341; 0.05 µg for SET1B, Active Motif #81342; 0.3 µg for MLL1, Active Morif #31423; 0.45 µg for MLL2, Active Motif #31499; 0.3 µg for MLL3, Active Motif #31478; 0.2 µg for MLL4, Active Motif #31498; 0.3 µg for G9a-SET 913-1193, Active Motif #31425; 0.1 µg for DOT1L 1-416, Active Motif #31474; 0.1 µg for SET1A, Reaction Biology, HMT-15-717; 0.05 µg for SET1B, Reaction Biology HMT-15-117), SAM (100 µM final) (NEB, B9003S), substrates (1 µg for histone H3.1, Active Motif #31894; 1 µg for histone H3.3, Active Motif #31894; 1 µg histone octamer, Active Motif #31470; 1.2 µg mononucleosome, Active Motif #81070; 0.6 µg polynucleosome, Active Motif #31466; H2BK120ub nucleosome, EpiCypher #16-0396), and various concentrations of compounds (dissolved in 2.5% DMSO) in the reaction buffer (50 mM Tris-HCl [pH 8.8], 20 mM KCl, 5 mM MgCl_2_, 0.5 mM dithiothreitol) was incubated at room temperature for 2 hours (SET1A, SET1B, DOT1L and G9a) or 3 hours (MLL1-4). SDS sample buffer was added to the reaction and samples were boiled at 95°C for 5 minutes prior to western blot for H3K4me1/2/3, H3K9me2, H3K79me1/2/3 and H3 to quantify the inhibitor IC_50_.

### MTase-GLO^TM^ assay

The MTase-Glo^TM^ assay was performed to evaluate the inhibitory effects of FCD-11 on the enzymatic activity of multiple histone methyltransferases. All assays were conducted in 384-well plates (OptiPlate, Revvity, #6007290).

To optimize assay conditions, two-fold serial dilutions of recombinant methyltransferases (SET1A, Active Motif #81341; SET1B, Active Motif #81342; MLL1, Active Motif #31423; MLL2, Active Motif #31499; MLL3, Active Motif #31478; and MLL4, Active Motif #31498) were prepared in 1× reaction buffer consisting of 20 mM Tris-HCl (pH 8.0), 50 mM NaCl, 1 mM EDTA, 3 mM MgCl₂, 0.1 mg/mL BSA, and 1 mM dithiothreitol (DTT). Histone H3.1 substrate (0.0375 μg/μL; Active Motif #31894) and S-adenosylmethionine (SAM; 10 μM final concentration; supplied in the MTase-Glo Methyltransferase Assay Kit, Promega #V7601) were maintained at constant concentrations. Reactions (8 μL total volume) were incubated at room temperature for 2 hours for SET1A and SET1B or 3 hours for MLL1–MLL4.

Following incubation, 2 μL of 5× MTase-Glo reagent (Promega #V7601) was added to each reaction, and plates were incubated for 30 minutes. Subsequently, 10 μL of MTase-Glo Detection Solution (Promega #V7601) was added and incubated for an additional 30 minutes before luminescence was measured using a TECAN Spark plate reader. Enzyme concentrations that produced signals above background and within the linear detection range were selected for subsequent experiments.

Using these optimized enzyme concentrations and assay conditions, additional substrates including histone H3 (1–25) amide peptide (Anaspec #AS-61703) and mononucleosomes (Active Motif #81070) were evaluated. In addition to COMPASS family methyltransferases, several non-COMPASS methyltransferases, including PRDM9 (residues 191–414; Active Motif #31510), SETDB1 (Active Motif #31452), G9a SET domain (residues 913–1193; Active Motif #31425), and DOT1L (residues 1–416; Active Motif #31474), were tested under the same assay conditions. These enzymes were incubated with substrates and SAM for 2 hours at room temperature.

Optimized assay conditions were subsequently used to determine the effects of FCD-11 on methyltransferase activity. Reactions (10 μL total volume) contained optimized concentrations of enzyme (SET1A, 0.01 μg/μL; SET1B, 0.015 μg/μL; MLL1, 0.05 μg/μL; MLL2, 0.05 μg/μL; MLL3, 0.025 μg/μL; MLL4, 0.025 μg/μL; G9a, 0.02 μg/μL; PRDM9, 0.005 μg/μL; SETDB1, 0.005 μg/μL; and DOT1L, 0.01 μg/μL), substrate (histone H3.1, 0.0375 μg/μL; mononucleosomes, 0.1 μg/μL; or histone H3 [1–25] amide peptide, 20 μM), SAM (5 μM final concentration), MTase-Glo reagent and a three-fold serial dilution of FCD-11 prepared in 2% DMSO. Reactions were incubated at room temperature for either 2 or 3 hours, depending on the enzyme, as described above.

Following incubation, 10 μL of MTase-Glo Detection Solution was added directly to each reaction, and plates were incubated for 1 hour at room temperature. Luminescence was then measured using a TECAN Spark plate reader. Dose-response curves were generated from normalized luminescence values, and absolute IC₅₀ values were calculated using nonlinear regression (log[inhibitor] versus response, three-parameter model) in GraphPad Prism software.

### Embryonic body (EB) formation

EBs were generated via a hanging drop method, in which 1200 cells were cultured in 23 µL of EB differentiation medium on the lid of 150-mm culture plates for 3 days. EB differentiation medium was used as previously described (*27*). Cells were pre-treated with DMSO or FCD-11 (2 µM) for 48 hours before seeding for EB generation, and treated with DMSO or FCD-11 (2 or 5 µM) during the formation of EB.

### ChIP-seq and analysis

ChIP-seq libraries were made using the KAPA HTP Biosystems Library Prep kit (Roche, KK8234) using Perkin Elmer Unique Dual Index Barcodes compatible with Illumina systems. DNA libraries were sequenced on an Illumina Nova-Seq 6000 instrument using paired-end 50 cycle reads. Reads were trimmed using Cutadapt v4.1 (*50*) with parameters --nextseq-trim=30 --minimum-length=20, then aligned to the mm10 mouse genome assembly or hg38 human genome assembly using Bowtie2 v2.2.6 (*51*) with parameters --sensitive --no-unal. For spikein, dm3 drosophila genome assembly was used. Peaks were called using MACS2 v2.1.0 (*52*) with parameters -q 0.05 --keep-dup auto --nomodel, then annotated using HOMER v4.1.1 (*53*). Protein coding genes detection and selection, TSS and gene body region selection, and coverage mapping were performed using in-house R and perl scripts. Occupancy plots and heatmaps were generated using deepTools v.3.5.1 (*54*). Subtraction tracks were made with UCSC’s Track Collection Builder function with Merge method set to “Subtract”.

### RNA-seq and analysis

RNA-seq samples were sequenced with Illumina NovaSeq technology on NovaSeq 6000, and output data were processed with bcl2fastq. Preprocessing was performed using CETO Toolbox (https://github.com/ebartom/NGSbartom). Sequence quality was assessed using FastQC v 0.11.2 (*55*), and quality trimming was done using Trimmomatic (*56*) with parameters TRAILING:30 MINLEN:20. RNA-seq reads were aligned to the GRCh38 genome for human and GRCm38 genome for mouse using STAR v.2.5.2 (*57*) and only uniquely mapped reads with a two-mismatch threshold were considered for downstream analysis and quantified to the gene level using HTSeq (58). Output BAM files were converted into bigWig track files to display coverage throughout the genome (in rpm). Gene count tables were constructed following Ensembl GRCh38 and GRCm38 (59). Differential gene expression analysis was performed using EdgeR (*60*). Genes with adjusted p values < 0.01 (the Benjamini and Hochberg method) were considered differentially expressed.

### Cellular thermal shift assay (CESTA)

Cellular Thermal Shift Assay (CETSA) was performed as previously described (*61*). Briefly, MCF7 cells were treated with DMSO or FCD-11 (20 µM) for 3 hours. Then, cells were harvested and re-suspended in PBS with protease inhibitor and heated in the thermocycler for 3 minutes at indicated temperature with two increments from 43°C to 55°C. After incubation for another 3 minutes at room temperature, cells were lysed in Triton lysis buffer and freeze-thawed for a total of 3 cycles in liquid nitrogen. After the final thaw, the samples were centrifuged at 20,000 g for 15 minutes at 4°C to remove cell debris containing unfolded proteins. The clear cell lysates were collected for Western blot.

### Xenograft studies

Six-week-old female athymic mice (nu/nu genotype, BALB/c background) were purchased from Envigo (Indianapolis, IN, USA) and housed under aseptic conditions. MDA-MB-468, MDA-MB-453, MDA-MB-231, and HCC-1954 cells were implanted into the mammary fat pad of athymic mice as previously described(*62*). Briefly, 4 x 10^6^ cells, in 0.4 ml of cell culture media with matrigel (BD Bioscience) at 1:1 ratio, were injected into the right mammary fat pad of mice under anesthetization by inhaled isoflurane. Mice were randomly assigned to vehicle (DMSO, n = 8) and FCD-11 treatment (10 mg/kg for 7 days, n = 8) groups when the size of tumors reached 100 mm^3^ (day 6 after implantation). Tumor size was measured with calipers on alternate days and the mice were euthanized when the tumor size reached 1000 mm^3^.

### Statistics

All data were obtained from at least three repetitions and were expressed as mean ± SD, as indicated in the figure legends. Two-tailed Student’s *t* test was applied for statistical analysis between two groups. Kaplan-Meier survival analysis was performed for animal studies, and the difference between survival curves was analyzed by the log-rank (Mantel-Cox) test. All statistical analysis was performed using GraphPad Prism 9. *P* values of less than 0.05 were considered significant.

## Study approval

All animal protocols were approved by the Institutional Animal Care and Use Committee at University of Alabama at Birmingham.

## Funding

National Institutes of Health grant R35-CA197569 (A.S.)

Liz and Eric Lefkofsky Innovation Research Awards (A.S.)

Epigenetic Therapies - New Approaches P50CA254897 Career Enhancement Program (Z.Z.)

## Author contributions

Conceptualization: Z.Z., R.K.M., A.S.

Methodology: Z.Z., M.K, L.S.

Investigation: Z.Z., M.K, L.S., R.L., C.B., E.U., K.M., J.W.

Visualization: Z.Z., M.K, M.I.,R.K.M.

Supervision: Z.Z., S.A.B., R.H., A.S.

Writing-original draft: Z.Z., M.K., S.G.

Writing-review & editing: Z.Z., S.G., A.S.

We also thank Dr. Yohhei Takahashi in collaboration with Dr. Rama Mishra for the initial in vitro characterization of COMPASS disruptor compounds and contributions for figure S2 and S4. We thank Cassandra N. Philips, Jordan D. John and Madhurima Das for technical support.

## Data and materials availability

Next-generation sequencing data are deposited at Gene Expression Omnibus (GEO) with the accession number GSE272792.

## Supplemental information

**Fig. S1. In vitro H3K4 methylation assay optimization.** (**A**) Western blot comparison of protein components in SET1A, SET1B, MLL1-4 SET complexes purchased from Active Motif. Blots were probed for ASH2L, RbBP5, WDR5, and DPY30. (**B**) Activities of SETDB1, PRDM9, DOT1L and G9a were evaluated using three different substrates: histone H3.1, histone H3 peptide (1-25 a.a.), and H3.1 containing mononucleosomes. n=4. (**C**) Western blot comparison of in vitro COMPASS H3K4 methyltransferase activity in assays performed with H3.3-containing histone H3 or mononucleosome substrates. Blots were probed for H3K4me1, H3K4me2, H3K4me3, and histone H3. (**D**) Western blot comparison of in vitro H3K4me3 HMTase activity using H3.1-containing histone H3 or mononucleosome substrates, for SET1A and SET1B complexes purchased from Active Motif (AM) or Reaction Biology (RB). Blots were probed for H3K4me3 and histone H3. (**E**) Silver staining comparison of COMPASS complexes purchased from Active Motif and those purified from 293T cells in-house. For in-house purification, 293T cells infected with lentivirus expressing 3xFlag tagged SET domain of SET1A, SET1B, or MLL1-4 were harvested, lysed, and immunoprecipitated against M2 slurry beads. (**F**) Western blot analysis of COMPASS complex immunoprecipitates to confirm intact WRAD interaction. Blots were probed for the FLAG tag, ASH2L, RbBP5, WDR5, and DPY30.

**Fig. S2. In silico and biochemical screens to identify a potent SET1A/COMPASS inhibitor.** (**A**) Schematic of the small molecule screening strategy using 3D structure of nucleosome-bound MLL1. (**B**) Western blot analysis of the first round of SET1A/COMPASS H3K4me3 methyltransferase inhibition assays with 99 compounds (100 µM) that were initially identified in the in-silico screen. (**C-D**) Second round of screening including SET1A H3K4me3 methyltransferase inhibition assays (**C**) and G9a H3K9me2 methyltransferase inhibition assay (**D**) with 34 compounds (10 µM) selected from (**B**). The compounds M2, M58, and M62 are highlighted in red. (**E**) Chemical structures of M2, M58, M62, and FCD-11. The core scaffold structure is shown on the right. (**F**) Comparison of FCD-11 and M62 IC_50_ in the SET1A H3K4me3 methyltransferase inhibition assay.

**Fig. S3. FCD-11 is a first-in-class COMPASS inhibitor.** (**A-D**) FCD-11 inhibition of methyltransferase activity in MTase-Glo^TM^ assays using histone H3 peptides (aa 1-25) (**A-B**) or mononucleosomes (**C-D**) as substrates. FCD-11 inhibition curve and absolute IC_50_ values are shown. G9a, PRDM9 and SETB1 were used as negative controls for assays using histone H3 peptides as substrates. DOT1L was used as a negative control for assays using mononucleosomes as substrates. (**E-F**) Western blot comparison of FCD-11 inhibition of in vitro SET1A (**E**) and MLL1 (**F**) HMTase activity in reactions using H2B K120 ubiquitinated mononucleosomes (H2BUb-Nuc) or unmodified mononucleosomes (Nuc) as substrates. Blots were probed for H3K4me1/2/3 and total H3 (loading control). (**G**) Western blot analysis of the histone methyltransferase (HMTase) inhibition assays for SET1A, SET1B, and MLL1-4/COMPASS; for G9a; and for DOT1L. Histone H3 was used as the substrate for all reactions except for those with MLL1 and DOT1L, where mononucleosomes were used. Increasing concentrations of FCD-11 were added to the reactions with DMSO as vehicle. Western blots were probed for H3K4me1/2/3 (for COMPASS HMTases), H3K9me2 (for G9a), and for H3K79me1/2/3 (for DOT1L), with total H3 as the loading control.

**Fig. S4. Structure-activity relationship (SAR) analysis of FCD-11.** (**A**) Western blot analysis of SET1A/COMPASS H3K4me3 methyltransferase inhibition assays for FCD-11 and 18 analogues. Q15, Q16, and Q17 were selected from this first round. (**B**) Western blot analysis of the second round of H3K4me3 methyltransferase inhibition assays, in which FCD-11, Q15, Q16, and Q17 were titrated from 0.04 to 10 µM. (**C**) Quantification of H3K4me3 band intensities in (**B**). (**D**) Chemical structure comparison of FCD-11 with Q15, Q16, and Q17.

**Fig. S5. FCD-11 treatment resembles functional SET1A/COMPASS loss in mESCs.** (**A**) Western blot demonstrating depletion of COMPASS methyltransferases in the clonal mESC cell lines. Blots for the Pol II subunit RPB1 (NTD) and the histone marks H3K4me1/2/3 and H3K27me3 are also shown. β-tubulin and total H3 were used as the loading controls. (**B**) Principal component analysis (PCA) of RNA-seq in cell lines with compounding COMPASS KO. ntop = 500. SET1AΔSET (A), SET1BKO (B), MLL1KO (M1), MLL2KO (M2), SET1AΔSET/SET1BKO (AB), SET1AΔSET/SET1BKO/MLL1KO (AB1), SET1AΔSET/SET1BKO/MLL2ΔSET (AB2), MLL3KO (M3), MLL4KO (M4), MLL3/MLL4KO (M34). (**C**) Hierarchical clustering heatmap of differential gene expression in the above cell lines. Genes with significantly differential expression in SET1AΔSET/SET1BKO/MLL2ΔSET versus WT cells were included in the analysis (excluding MLL3KO, MLL4KO, and MLL3/MLL4KO). (**D**) Histone H3K4me1/2/3, H3K36me2/3 H3k79me1/2/3, H3K9me2/3, and H3K27me2/3 levels were examined by western blot after FCD-11 treatment for 72 hours in WT mESCs (0, 0.5, 1, and 2 µM). (**E**) Principal component analysis (PCA) of H3K4me3 ChIP-seq in the above cell lines in **A-B**. (**F**) Feature distribution of SET1A, MLL1 and MLL2 chromatin binding. SET1A, MLL1 and MLL2 ChIP-seq were obtained from GSE200120(*28*). (**G**) H3K4me3 changes with FCD-11 and its analogues on the kmeans clustering defined in Fig. 3A. (**H**) ChIP-seq of histone marks with DMSO or 2 µM FCD-11 treatment for 48 hours. Log2FC heatmap was shown with the same clustering as in Fig. 3A. (**I-J**) Boxplot showing the quantification of differential expression in cluster1 and cluster2 genes, for SET1AΔSET versus WT (**I**) and for FCD-11 versus DMSO (**J**). (**K**) Embryonic body (EB) formation and EB size quantification in WT and SET1AΔSET cells and WT cells treated with FCD-11 at indicated concentrations.

**Fig. S6. FCD-11 decreases cell viability, diminishes SET1A/COMPASS-deposited H3K4me3 peaks, and selectively engages SET1A/COMPASS in breast cancer cell lines.** (**A**) One-dose (10 µM FCD-11) mean growth plot from the NCI-60 cell line viability screen for FCD-11 treatment performed by the Developmental Therapeutics Program. (**B**) FCD-11 CellTiter-Glo® luminescent cell viability assay for a panel of cell lines treated with increasing concentrations of FCD-11. IC_50_ values of FCD-11 in these cell lines are summarized in the table. (**C**) Correlation between FCD-11 IC_50_ and CRISPR score (dependency score) for other the COMPASS HMTases, for COMPASS WRAD components (WDR5, RBBP5, ASH2L, DPY30), for SET1A/B-COMPASS-specific subunits (HCFC1, CXXC1, WDR82), and for other non-COMPASS HMTases (EHMT2, EZH2, PRDM9, DOT1L, SETD2, and SETDB1). (**D**) Differential ChIP-seq peak analysis showing H3K4me3 peaks that are increased or decreased when treated with 2.5 µM FCD-11 relative to DMSO. 2463 upregulated peaks, adj.p < 0.01. 376 downregulated peaks, adj.p < 0.01.

