## supplemental figure 1-6 for "FCD-11: A First-in-Class COMPASS Inhibitor in Cancer Therapeutics"

**Fig. S1. H3K4 methylation in vitro assay optimization.**

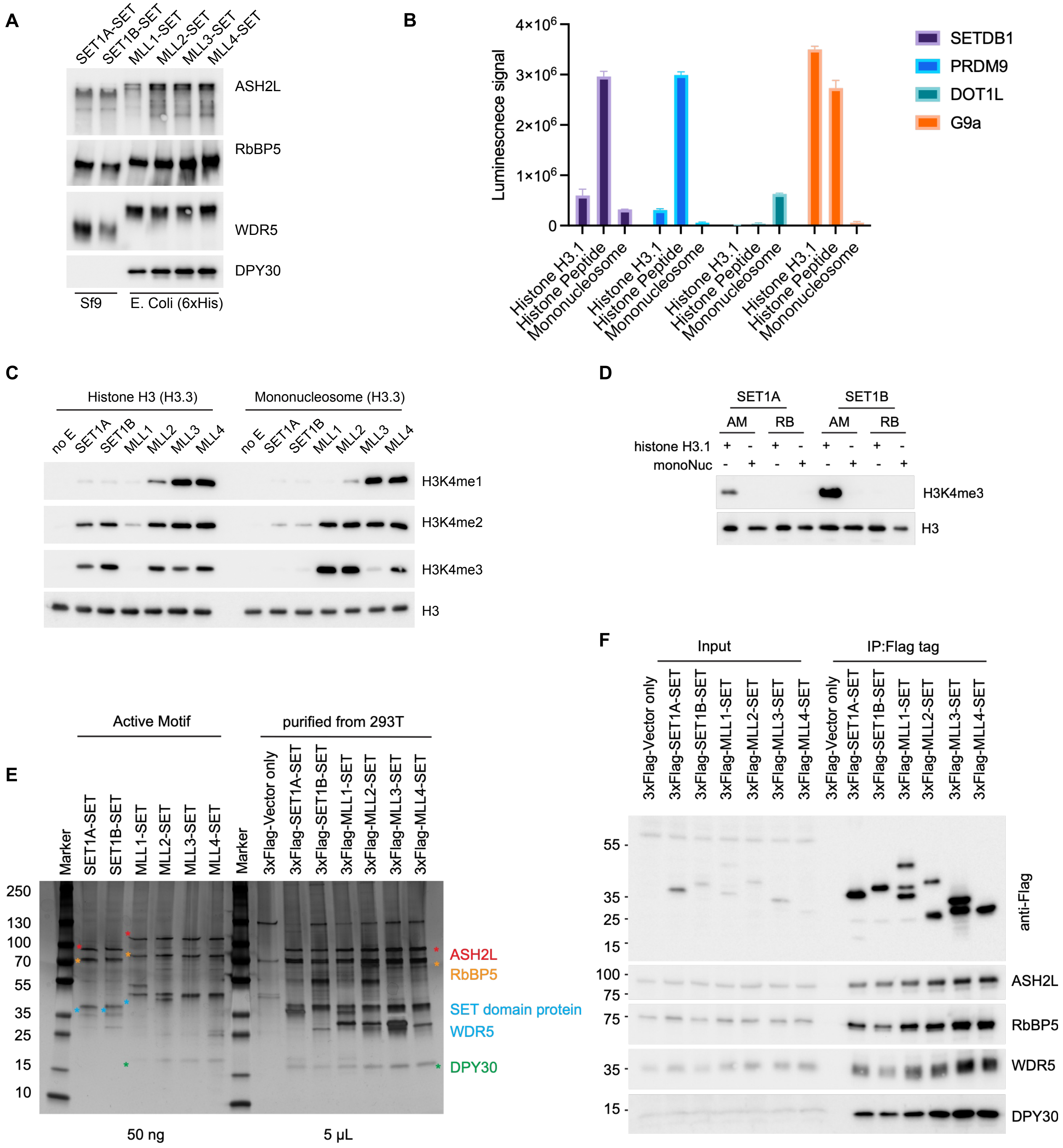

**Fig. S2. In silico and biochemical screens to identify a potent SET1A/COMPASS inhibitor.**

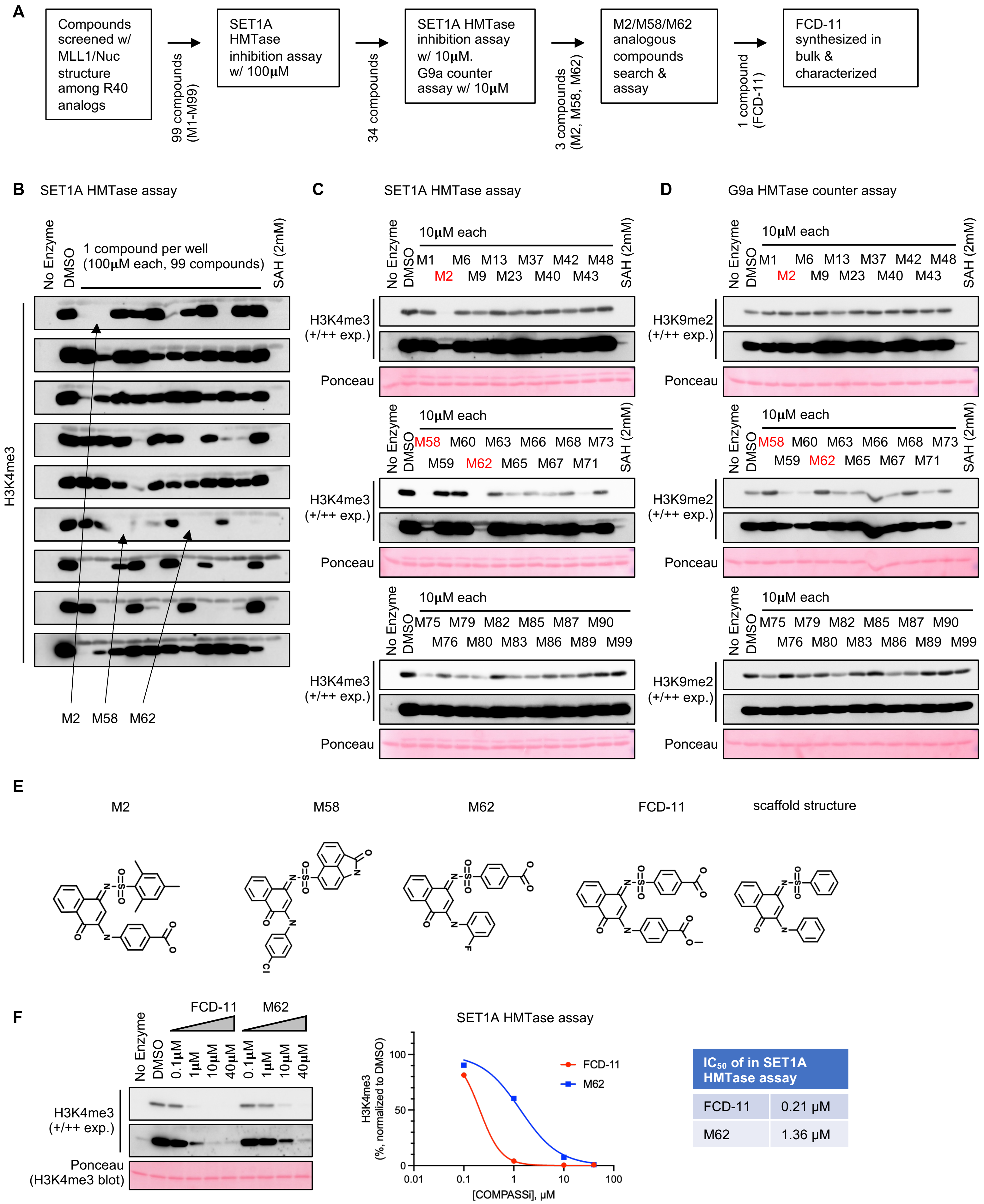

#### Fig. S3. FCD-11 is a first-in-class COMPASS inhibitor.

A

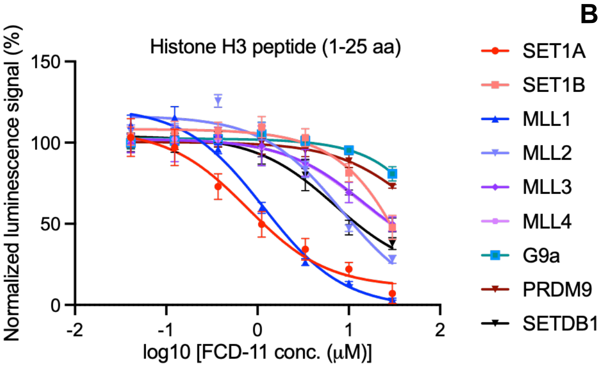


B

| Enzymes | $IC_{50}$ (µM) |
|---|---|
| SET1A | 1.11 |
| SET1B | 27.9 |
| MLL1 | 1.6 |
| MLL2 | 10.7 |
| MLL3 | 27.7 |
| MLL4 | 27.1 |
| G9a | >30 |
| PRDM9 | >30 |
| SETDB1 | 11.7 |

C

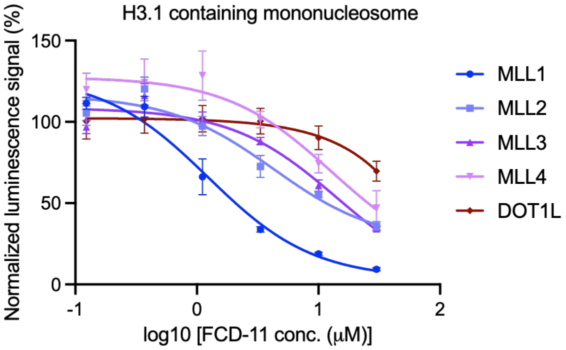


D

| Enzymes | $IC_{50}$ (µM) |
|---|---|
| MLL1 | 2.01 |
| MLL2 | 10.96 |
| MLL3 | 15.13 |
| MLL4 | 24.7 |
| DOT1L | >30 |

E

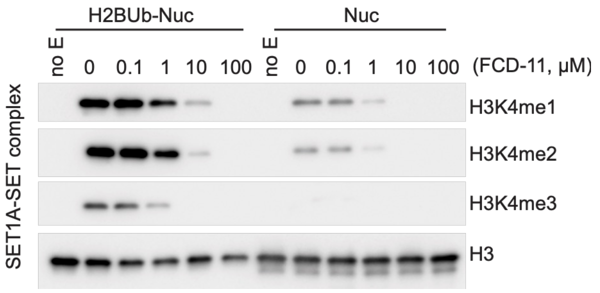


F

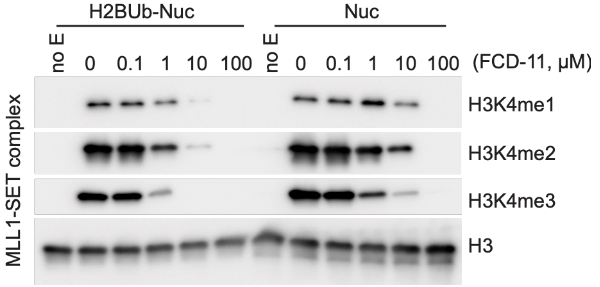


G

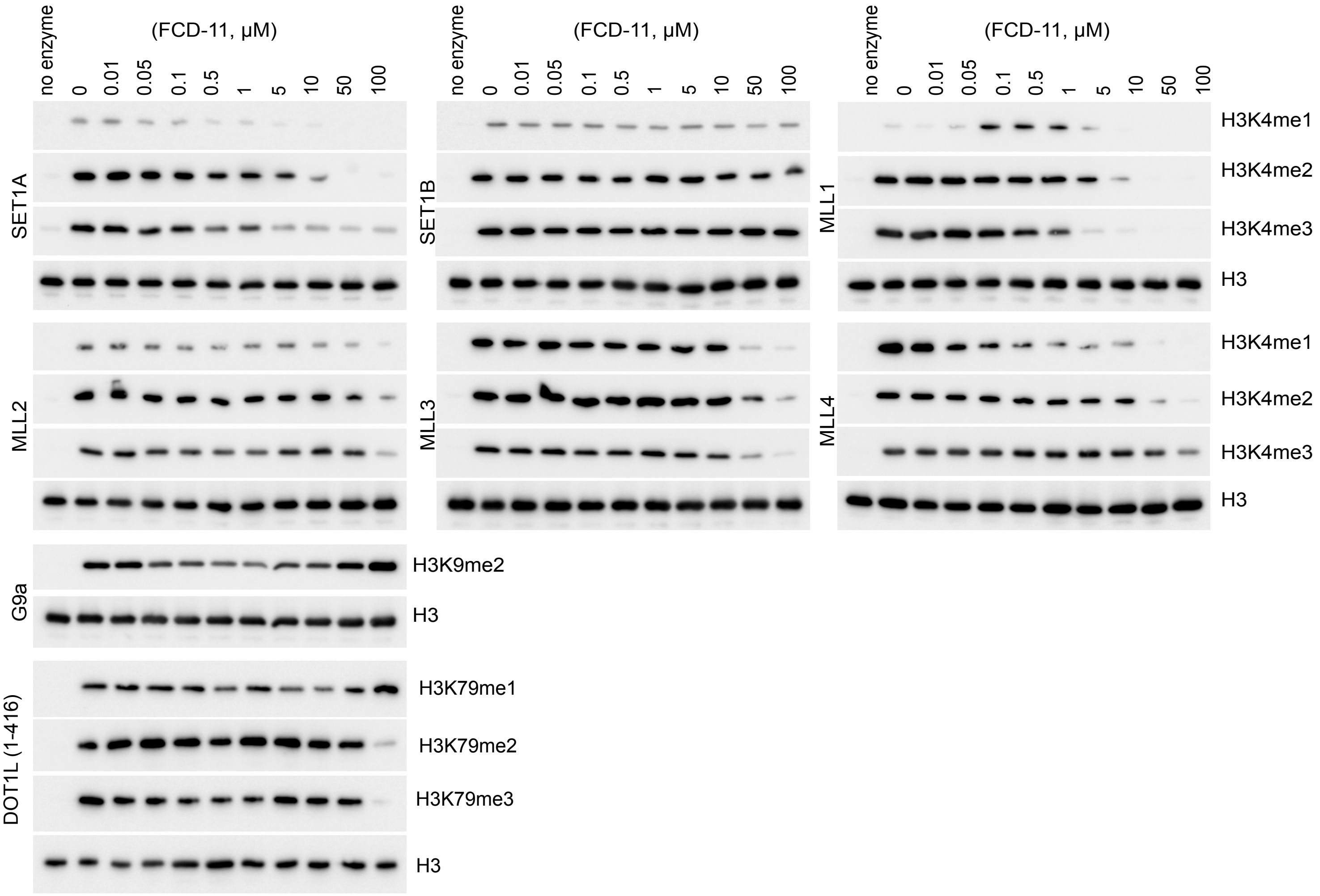

**Fig. S4. Structure-activity relationship (SAR) analysis of FCD-11.**

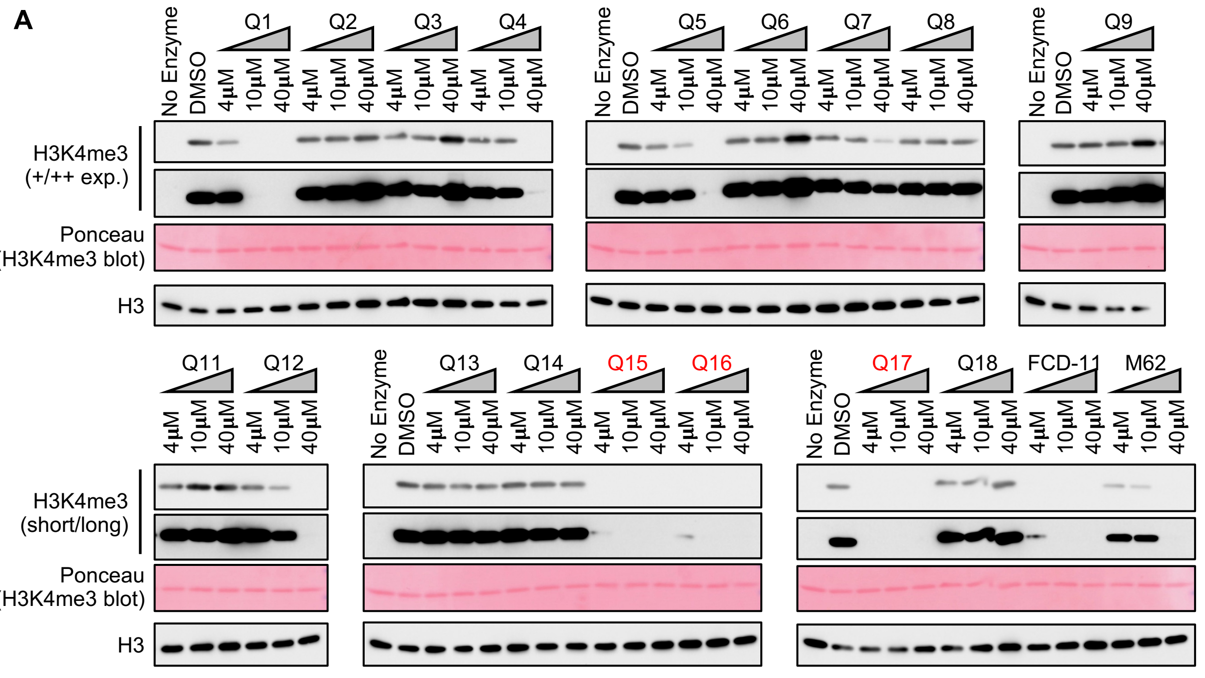


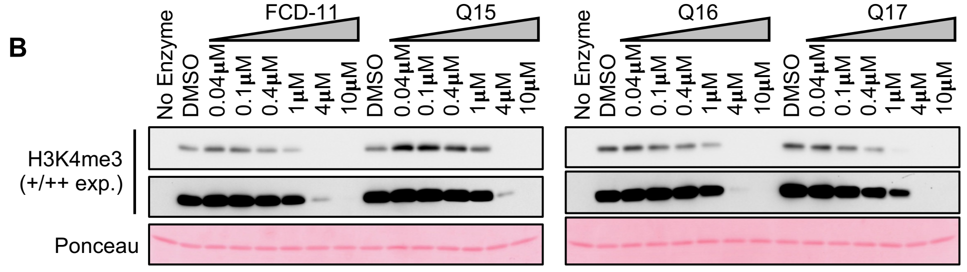


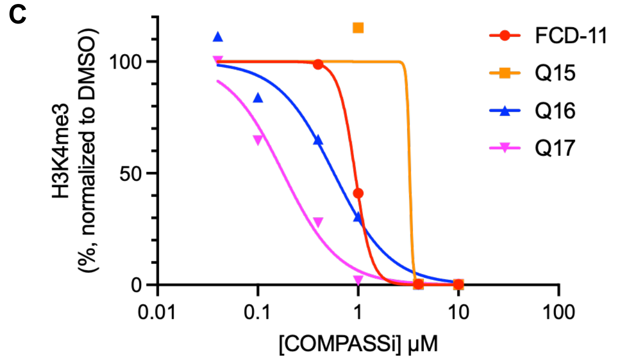


| IC$_{50}$ of FCD-11 analogues in SET1A HMTase inhibition assay | |
|---|---|
| FCD-11 | 0.93 μM |
| Q15 | 3.2 μM |
| Q16 | 0.57 μM |
| Q17 | 0.18 μM |

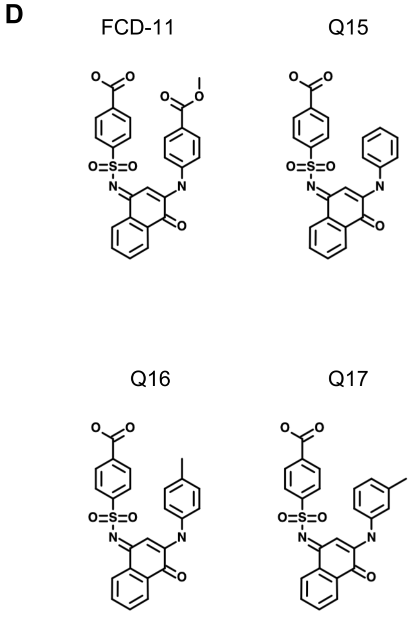

### Fig. S5. FCD-11 treatment most closely resembles functional SET1A/COMPASS loss in mESCs.

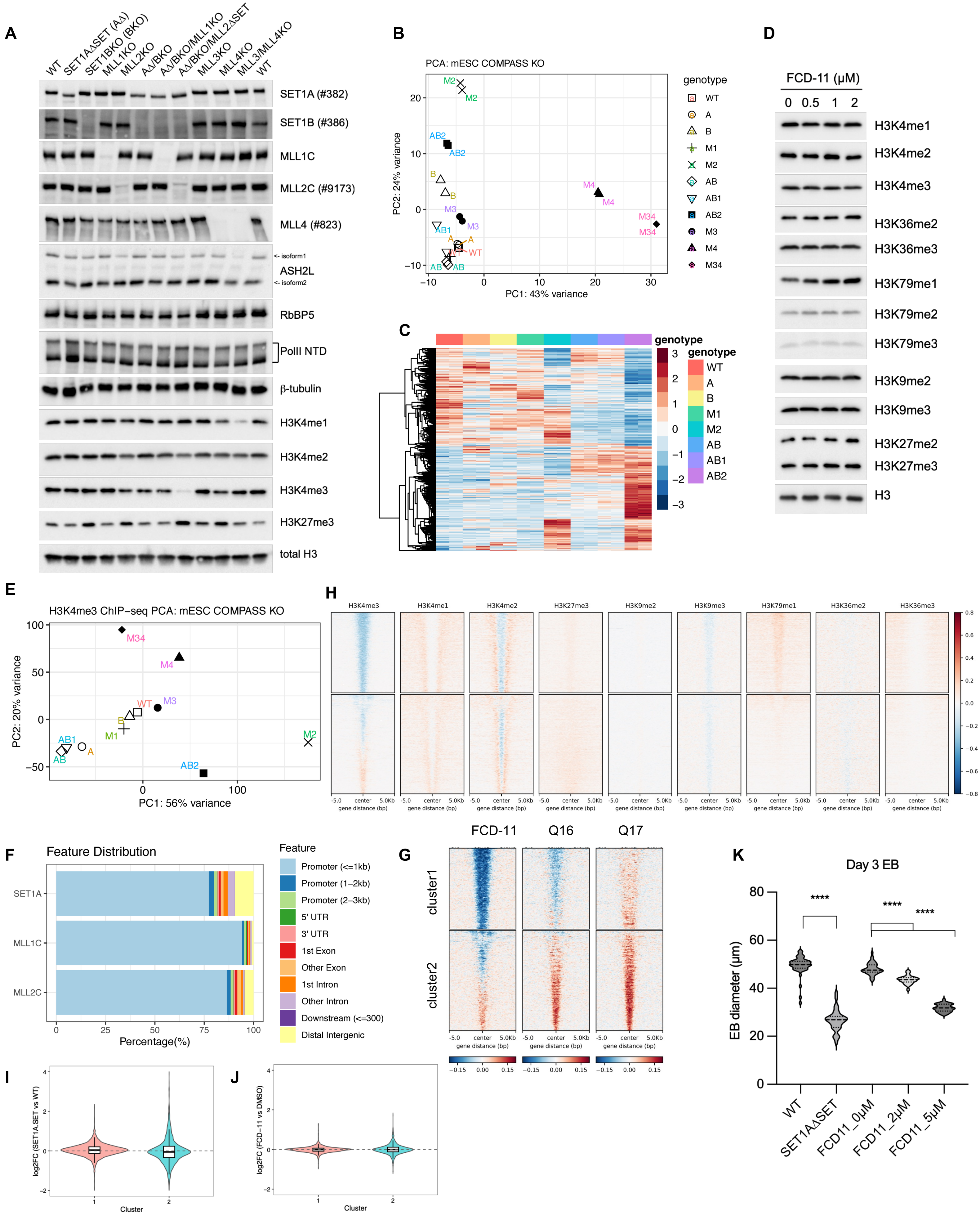

**Fig. S6. FCD-11 decreases cell viability, diminishes SET1A/COMPASS-deposited H3K4me3 peaks, and engages SET1A/COMPASS in SET1A-dependent breast cancer cell lines.**

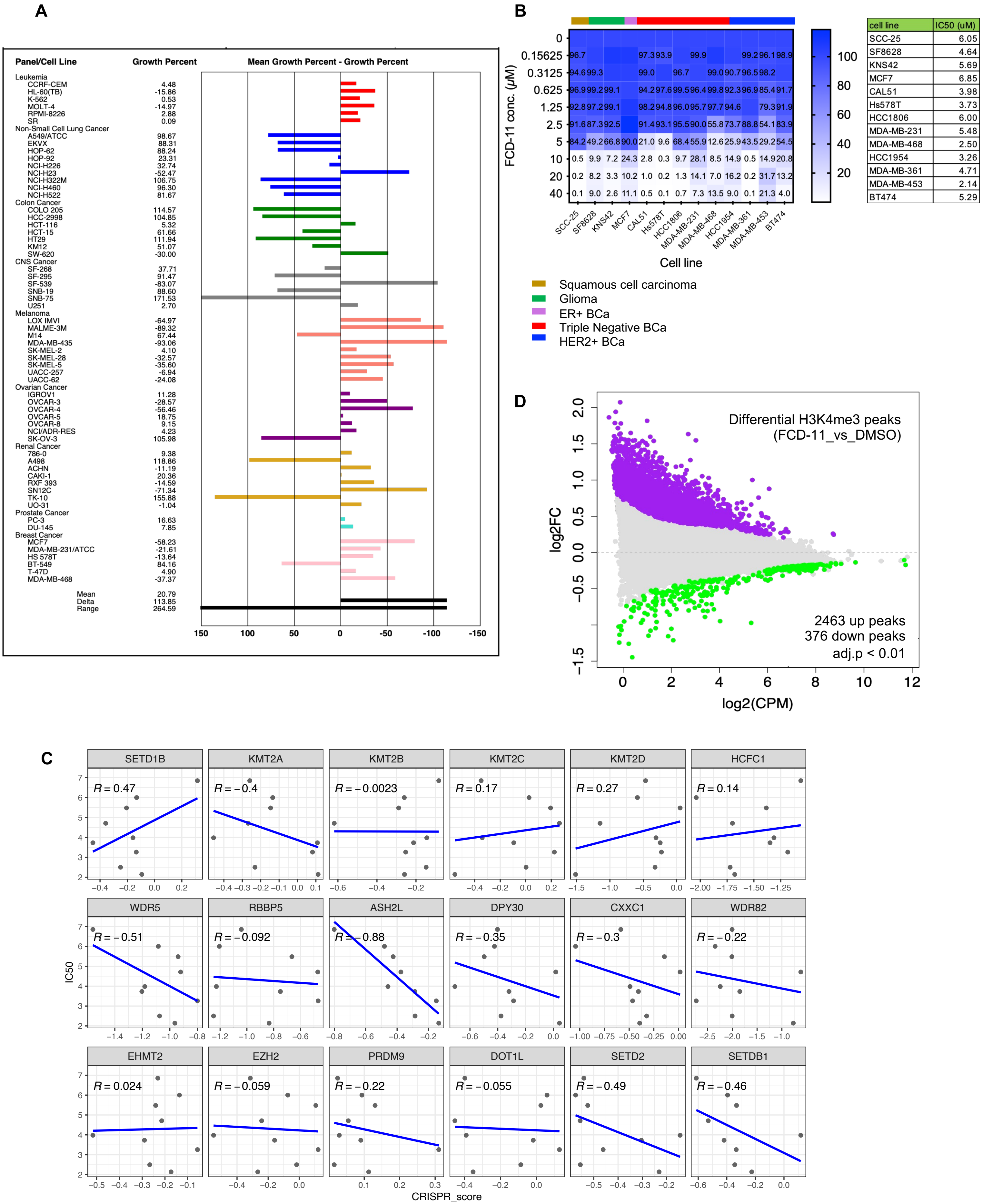


| cell line | IC50 (uM) |
|---|---|
| SCC-25 | 6.05 |
| SF8628 | 4.64 |
| KNS42 | 5.69 |
| MCF7 | 6.85 |
| CAL51 | 3.98 |
| Hs578T | 3.73 |
| HCC1806 | 6.00 |
| MDA-MB-231 | 5.48 |
| MDA-MB-468 | 2.50 |
| HCC1954 | 3.26 |
| MDA-MB-361 | 4.71 |
| MDA-MB-453 | 2.14 |
| BT474 | 5.29 |
